# Computational design and experimental characterization of mini-protein binders targeting Nipah, Langya and Measles virus receptor-binding domains

**DOI:** 10.64898/2025.12.20.695652

**Authors:** Dominic Rieger, Carolin Rüdiger, Antonia Sophia Peter, Phillip Schlegel, Anna Nobis, Hannes Junker, Lena Kiesewetter, Max Beining, Lorenz Beckmann, Johannes Klier, Nichakorn Pipatpadungsin, Robert Stass, Thomas A. Bowden, Sandra Diederich, Jens Meiler, Clara T. Schoeder

## Abstract

A lack of reagents represents a major bottleneck in pandemic preparedness and rapid vaccine development. It is therefore important to enable the design of reagents for use in the treatment and diagnosis of emerging viral diseases. Ideally, the design and identification platform is fast, can be performed by testing only a small number of candidates and enables a generally applicable strategy. In this study, we assessed the ability of recently developed computational protein design tools to establish such a workflow for validating paramyxovirus receptor-binding protein *de novo* binders as such reagents. The family *Paramyxoviridae* includes various members that cause severe disease and exhibit re-occurring zoonotic spillover events, with documented human infections over the past decades. We successfully designed, identified, and characterized mini-proteins targeting the receptor binding proteins of Nipah virus, Langya virus, and Measles virus while screening as few as 10-16 designs per target. The resulting functional binders have moderate to low nanomolar affinities and display high on-target specificity. We further showed that our most promising Nipah virus receptor-binding protein binder is able to inhibit human receptor binding *in vitro* and competes for an epitope that overlaps with that of the neutralizing antibody HENV-117. However, despite these promising results, this Nipah binder is only weakly neutralizing, preventing therapeutic applications. Nevertheless, we established a platform, applicable to rapidly generate diagnostically relevant proteins from only a small number of candidates, and developed novel reagents for the *Paramyxoviridae* family.

## Introduction

Viruses from the *Paramyxoviridae* family, such as Measles virus (MeV) or Mumps virus, continue to cause severe outbreaks, particularly in populations with sub-optimal immunization coverage.^1–3^ With zoonotic spillover events originating from bat or rodent reservoirs, emerging pathogenic paramyxoviruses pose a threat to human health . For example, the henipaviruses, Nipah virus (NiV) and Hendra virus (HeV), cause severe respiratory and neurological symptoms with fatality rates up to 70%.^4–7^ Further, a new henipa-like virus, termed Langya (LayV), has been identified that originates from shrews and has the potential to infect humans.^8–10^ The potential for zoonotic paramyxoviruses to emerge from host reservoirs and cause disease has catalyzed a wave of research to better understand these pathogens.^6,11–13^ Indeed, although more recently discovered members have not caused as severe outbreaks and symptoms as NiV or HeV, these events mandate the development of vaccines, antiviral drugs, diagnostics, and basic research with regard to virus transmission, as pandemic preparedness measure.^14–18^

A common feature of all paramyxoviruses is the presence of two viral surface proteins, a receptor-binding protein (RBP) and fusion glycoprotein (F).^19^ These glycoproteins work in concert to negotiate viral entry into a host cell.^20,21^ The RBP can be divided according into receptor-binding functionality, where hemagglutinin neuraminidase (HN) RBPs bind and hydrolyze sialic acid, hemagglutinin (H) RBPs are presented by morbilliviruses and bind signaling lymphocyte activation molecule (SLAM) and nectin-4, and attachment (G) glycoproteins bind to other proteinaceous receptors.^22,23^ The RBP consists of a C-terminal cytoplasmic tail, a transmembrane domain, and an extracellular region consisting of a tetramer-inducing stalk region and C-terminal six-bladed β-propeller, referred as receptor-binding domain (RBD) (Supplementary Fig. 1).^24–26^ It has been shown that receptor binding of G/H/HN triggers the fusion protein F to mediate host and virus membrane fusion to release the viral genome into the cells and thus, infect cells.^21,27,28^ As seen with several neutralizing antibodies, blocking the interaction of the RBP with the receptor can protect against infection.^29,30^ Therefore, a common strategy for immunization is vaccination with the respective paramyxovirus RBP.^31,32^ However, it has also been shown that for some RBPs, such as LayV G, stabilizing mutations are required to allow for expression as soluble glycoprotein.^25^ It is critical for vaccine development, but also for proteinogenic diagnostics and antiviral therapeutics to discover antibodies or other protein-binding domains that specifically target the protein of interest, allowing for probing of antigen identity, quantity, or structural integrity.^33–35^

Traditionally, antibody discovery is performed to identify protein-binding domains, resulting in highly potent and often neutralizing monoclonal antibodies.^30,36^ Human samples from survivors are often scarce and hard to obtain. Therefore, these identification strategies can be time consuming and require extensive experimental costs.^37^ Within the past years, the emergence of artificial intelligence in protein design has enabled new strategies that have high success rates and potentially circumvent the requirement of sample acquisition or large library high-throughput screenings.^38–41^ These *in silico* design approaches been proven to be effective in the development of antiviral or antimicrobial mini-proteins by targeting surface-proteins, e.g. for the influenza hemagglutinin or MERS-CoV spike glycoprotein RBD.^39,42,43^ The typical properties of designed mini-protein binders, such as thermostability, high-yield production, target specificity, and high affinity, make it a promising alternative to monoclonal antibodies, especially in diagnostics and also as biotherapeutics.^42,44–46^

Here, we sought to assess the success and the generalizability of computational protein design to generate mini-protein binders for paramyxovirus receptor-binding proteins. We developed an *in silico* workflow with subsequent *in vitro* validation to identify binders against the well-studied NiV G protein, but also against the G protein of the newly emerged LayV. To probe for transferability of our platform to more distantly related RBPs, we chose MeV H as additional target. Our assessment focused on the success of binder design for each target, and the characterized properties of designed mini-proteins, respectively. We obtained binders for all three targets and thoroughly characterized their binding, specificity, stability and structure, including two *apo* crystal structures of designed mini-proteins. Initial designs were optimized through computational and rational engineering, however, no single approach was superior across all three targets. Finally, we validated one NiV G binder further for antibody and host receptor competition, as well as neutralization of infectious NiV. Overall, our results indicate that computational protein design is a fast and efficient strategy to obtain specifically targeting binders for paramyxovirus RBPs using low throughput experimental screening technologies.

## Results

### Computational design of mini-proteins targeting paramyxovirus RBPs

We chose three different members of the *Paramyxoviridae* family, namely NiV, LayV and MeV, to investigate the design of mini-protein binders against their respective RBD. Initially, RFdiffusion and ProteinMPNN were applied as generative models.^39,47^ After its release, BindCraft was utilized as well.^38^ Despite the same fold of all paramyxovirus RBDs, the desired target epitope of the host receptor interaction can differ significantly.^48^ Additionally, the glycosylation pattern and oligomerization of the G/H protein alters within the three target proteins of interest, NiV G, LayV G and MeV H (Supplementary Fig.1).^49,50^ Therefore, each target was analyzed separately to specify suitable residues in the host receptor interface that served as hotspots for *in silico* binder design. For NiV G and MeV H, the known receptor binding site with ephrin B2/B3 and SLAM/nectin-4 was targeted, respectively (Fig. 1a, Supplementary Fig.1a, c). ^51,52^ The co-crystal structures of receptor and RBD were used for the design rationale but binders were generated against the unbound structures. Generally, the design and selection of binders was solely performed using the RBD of the G/H protein which is frequently used in the research of paramyxoviruses as the receptor tropism-dictating region.^26,53–56^ It was favored over the full ectodomain to avoid avidity in affinity measurements and the requirement of stabilizing the RBP tetramer.^25^ However, the RBD of MeV H is expected to be dimeric in solution (Supplementary Fig.1c).

**Fig. 1.**
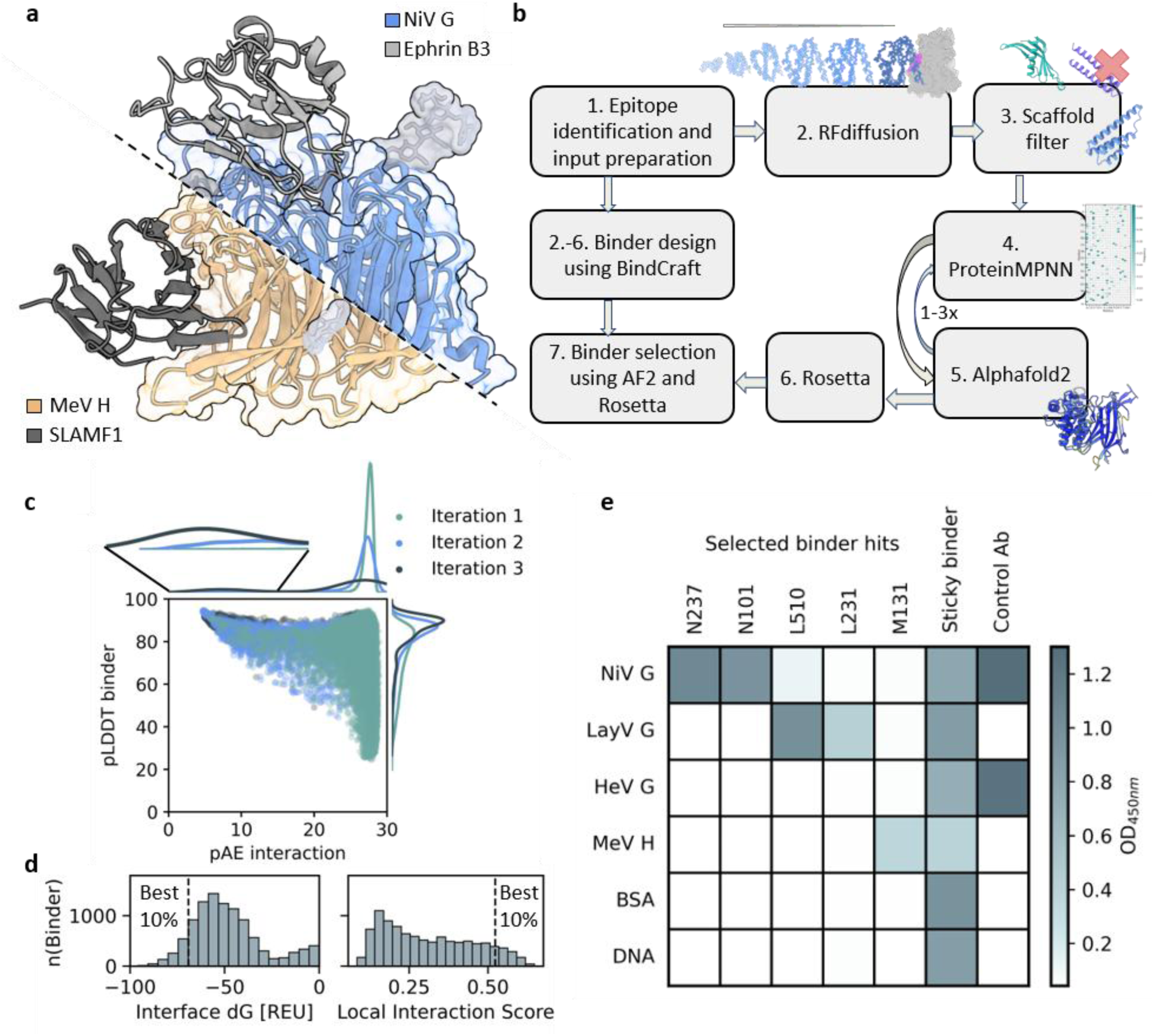
Computational design of *de novo* binder targeting NiV G, LayV G and MeV H RBD. **a**, Comparison of Ephrin B3-bound NiV G (PDB: 3D12) and SLAMF1-bound MeV H (PDB: 3ALZ). **b**, Schematic overview of *de novo* binder design workflow using BindCraft or a modified pipeline inspired by Bennett et al. ”dl_binder_pipeline”, including an additional filter step after backbone generation and iterative cycles of sequence design and structure prediction.^38,39,47,78^ **c**, Example library of binders. Predicted local distance difference test (pLDDT) of the binder and predicted aligned error of the interaction (pAE interaction) are shown colored by iteration of sequence design and structure prediction. On top and on the right of the scatter plot, densities of predicted binder confidences are shown as kde plot. **d**, Example of final filtering, based on Rosetta scores (e.g. Interface dG) and structure prediction confidences (e.g. AF2ig Local Interaction Score).^63^ **d**, Summarized enzyme-linked immunosorbent assay (ELISA) results of binder design targeting NiV, LayV and MeV RBD using selected examples at a concentration of 300 nM. Binding in the ELISA is shown measured using optical density at 450 nm (OD_450nm_).

The selected residues for NiV G (PDB 3D11) F358, Y518, and I588 are located in the loops on top of the β-propeller domain (Supplementary Fig. 1a). This selection resulted in binders reaching into the pocket as observed for ephrin B3 (Fig.1a).^24^ Residues Y541 and F522 of MeV H (PDB 2ZB6) are located on the β-sheet that interacts with SLAM.^57^ This allowed design to interact via hydrogen bonds with the outermost β-strand backbone (Supplementary Fig. 1c), a design strategy commonly known as β-pairing.^58,59^ As LayV belongs to henipa-like viruses, the same epitope as for NiV G was targeted, despite the fact that LayV G does not bind NiV/HeV-cellular-receptors.^60^ The structure, including residues P311, M313, F403, was remodeled using Colabfold due to unresolved loops in the CryoEM structure and loop crystal contacts in the crystal structure.^25,61^ Noteably, the LayV G input structure for design, predicted with templates, displayed an epitope backbone RMSD of 0.64-0.89 Å to predicted AF3 models, despite training data did not include experimental LayV G structures. This demonstrates the ability to design binders against the RBD of new emerging paramyxoviruses in terms of accurate structure predictions for the design input preparation.^62^

In all cases, epitope-adjacent glycosylation sites were set as forbidden contacts and generated RFdiffusion backbones in proximity to these were filtered out prior to sequence generation. Additionally, designs with helix-loop-helix topology and inaccessible C-terminus, in regard to Fc fusion, were removed in the scaffold filtering (Fig. 1b). Within the RFdiffusion-based binder design workflow, the iterative sequence design and complex prediction, led to improved confidences in structure prediction with Alphafold2 initial guess (AF2ig) (Fig. 1c).^39^ Especially in the last iteration, the high confidence binders converged towards a few scaffolds with relatively low sequence diversity, indicating enrichment based on AF2ig predictions. However, only unique RFdiffusion backbone scaffolds were selected for the experimental validation. Rank-based cutoffs were applied based on prediction confidences and biophysically-based calculated metrics. The library was filtered to include only those designs that fell within the top-scoring fraction for several criteria (Fig. 1d). Hence, the average scores of the final selection also differed on the target protein input. For example, the pAE-based local interaction score (LIS) for selected binders average for NiV G and MeVH at 0.57, and LayV G at 0.65.^63^ The final selection of designs was conducted by visual inspection of candidates that satisfied the predefined thresholds.

### Identification of functional RBD binders

We designated NiV G, LayV G, and MeV H binders with the prefixes N, L, and M, respectively. In total, 10 N-binders, 16 L-binders (including 6 BindCraft designs), and 15 M-binders (including 5 BindCraft designs) were selected for recombinant expression in Expi293 mammalian cells. Binders with a C-terminal Fc tag were screened by enzyme-linked immunosorbent assay (ELISA) and biolayer interferometry (BLI) using either purified or unpurified mini-proteins (Supplementary Fig. 2). The majority of designed proteins was well expressed, and binding was identified for all three target proteins (Fig. 1e, Supplementary Fig. 2). Binding was considered specific when the response for the on-target was significantly higher than for respective measured off-targets. We further validated binders through determination of their respective K_D,_ using BLI (Fig. 2, Supplementary Fig. 3a-e). The low-throughput screening revealed two binders within the nanomolar range for each target (Fig. 2, Supplementary Fig. 3c). Additionally, low affinity binders and unspecific binders, that bound DNA and BSA, were identified, indicating substantial off-target binding or stickiness (Supplementary Fig. 2, 3g).

**Fig. 2.**
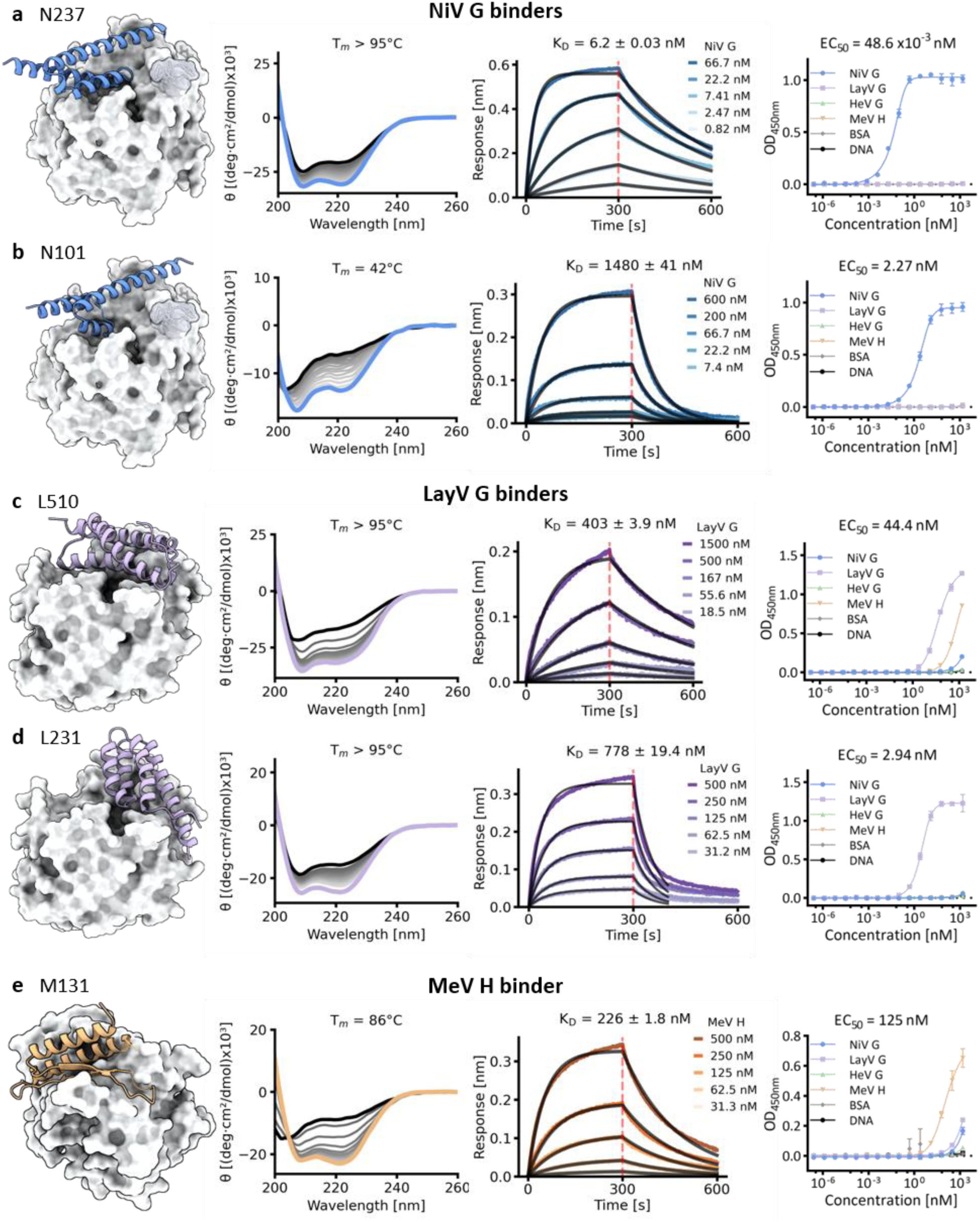
Characterization of NiV, LayV and MeV RBD binders. **a-e**, From left to right: Predicted model of binder-target-complex using AF2ig. Circular dichroism measurements including temperature ramp to observe thermal unfolding with the melting temperature (T_M_) calculated from the ellipticity at 222 nM as function of the temperature. BLI binding affinity measurements, including processed data and fit. ELISA EC_50_ determination against a panel of paramyxovirus RBDs and unrelated controls. EC_50_ is reported for the desired target, respectively. Characterized binders include **a**, N237, **b**, N101, **c**, L510, **d**, L231, **e**, M131.

### Characterization of binder hits revealed designs with different biophysical properties

We further analyzed the binder candidates N237, N101, L231, M131, designed with RFdiffusion/ProteinMPNN, and L510, designed with BindCraft, in regard to their biophysical properties. For all five proteins, the CD measurement of tagless variants recombinantly expressed from *E. coli*, displayed helical content, partially confirming the correct secondary structure of the designs (Fig. 2). Whereas NiV G and LayV G binders were helical bundles (Fig. 2a-d), the indented β-pairing target region of MeV H resulted in a mixed α/β scaffold that additionally contained an unstructured loop interacting with the target protein, as predicted (Fig. 2e). Except for N101, all characterized binders displayed a moderate to high thermostability (Fig. 2). Generally, N101 represented an exception as it was the only selected design with less than 75 residues, due to manual removal of an elongated, non-interacting N-terminal helix. It therefore possesses only a rather small protein interior (Fig. 2b). The expression with Fc tag in Expi293 cells and expression in *E. coli*, with tag removal during the purification, usually yielded in high amounts of pure protein (Supplementary Fig. 2, 4b). Only L510 showed minor expression in the mammalian expression system, but high expression from *E. coli* despite human codon optimization.

The highest affinity binder for each target was NiV G binder N237 with 6.2 nM, MeV H binder M131with 226 nM, and LayV G binder L510 with 403 nM (Fig. 2) as determined by BLI. Confirmatory K_D_ measurements with Isothermal calorimetry (ITC) resulted in 8.6 nM for N237, 975 nM for M131, and 48.2 nM for L510 (Supplementary Fig. 4c, d). Interestingly, except for N237, the measured affinities by BLI and ITC differed, which might be due to the altered constructs, Fc tag versus tagless, or different measurement types, kinetic versus equilibrium. Interestingly, within the same workflow we generated *de novo* binders that mediate their interaction solely through entropic contributions, and binders that displayed both enthalpic and entropic contributions (Supplementary Fig. 4d). In ELISA experiments, identified hits showed high specificity towards the desired target, with our most affine binder, N237, displaying only 3.3-fold weaker binding than the control antibody HENV-117 (Fig. 2, Supplementary Fig. 3f). Interestingly, L510 and M131 showed minor responses for the other paramyxovirus RBDs at high concentrations, but unlike unspecific binders, did not show binding to BSA or DNA (Fig. 2c, e, Supplementary Fig. 3g). The specificity profile of the binder was especially noteworthy for N101 and N237. The off-target panel included HeV G RBD which has a sequence identity of 77% and sequence similarity of 89% to NiV G (Supplementary Fig. 5a). Both RBPs interact with the same human receptors and several antibodies, such as HENV-117, have been shown to be cross-reactive (Supplementary Fig. 3f).^64,65^ In contrast, both NiV G binders target residues on NiV G adjacent to the receptor binding site that are distinct in both henipaviruses (Supplementary Fig. 5a). Interestingly, both binders contain identical interface residues in this region (Supplementary Fig. 5a). This demonstrates that the same interactions can be generated from different scaffolds. Taking the frequency of generated and well-scored interactions into account was beneficial for the selection of LayV G binders (Supplementary Fig. 6d). We sought to select 10 binders (RFdiffusion/ProteinMPNN-based) from the scored and filtered library, where five of them contained many frequently occurring protein-protein interaction patterns and five with equal scores and confidences but more unique interactions within the generated library. All three RFdiffusion/ProteinMPNN validated LayV G binders corresponded to the common interface interaction group (Supplementary Fig. 2b, c).

Despite the successful binder design for all desired targets, the affinities ranged from low to high nanomolarity. We hypothesized that at this point the binding strength would need to be improved for therapeutic use, as for example, potent antibodies are often in the sub-nanomolar range. We sought to evaluate various structure-guided affinity maturation strategies, excluding library-based high-throughput screenings. Here, we focused on either mutating single residues or regenerating the entire sequence based on the prediction of the functional scaffold (Supplementary Fig. 6a).

### Strategies for structure-based optimization of binders via point mutations

Three different computational approaches were chosen to predict beneficial point mutations in the predicted protein-protein interface of N237 (Supplementary Fig. 6c, e, f). In total, twelve mutations in N237, spanning over the entire interface (Fig. 3a), and one mutation in N101 were tested experimentally. Whereas almost all mutations in N237 displayed an effect on binding, only three mutations were beneficial (Fig. 3b, Supplementary Fig. 8). The two most promising mutations, located at the same position in the interface, S47, exhibited a 2-fold improvement. By increasing the sidechain size from serine to methionine or aspartic acid, newly forming interactions improve binding. N237_S47D, originating from a ProteinMPNN/AF2ig consensus sequence, displayed a K_D_ of 3.8 nM representing our overall most affine binder (Supplementary Fig. 8). Noteworthy, none of the approaches did generate any deleterious mutations, while each generated at least one that, although marginally, increased the affinity. The single point mutation introduced into N101 exhibited significant improvement from a K_D_ of 1480 nM to 252 nM, corresponding to an almost 6-fold improvement.

**Fig. 3.**
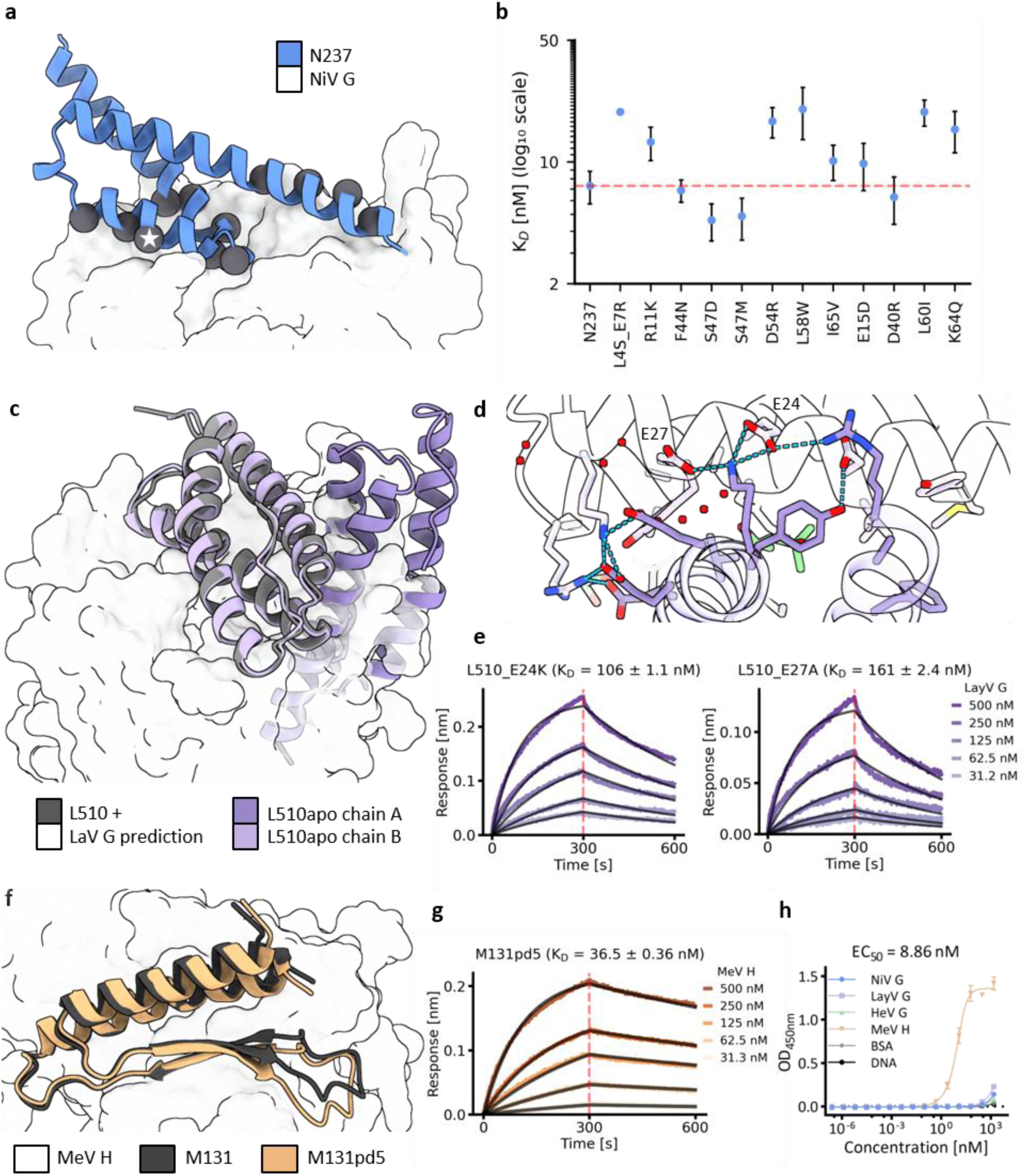
Structure-based strategies for the optimization of functional designs. **a**, Predicted N237-NiV G model with the position of the twelve individual point mutations marked as spheres. Position S47 is highlighted with a star. **b**, K_D_ values determined from BLI measurements are plotted as average from two biological replicates with standard deviation, except L4S_E7R. The K_D_ of the original N237 is shown as a red line. **c**, Superimposition of the L510-LayV G prediction with the crystal structure of L510 reveals a clashing dimer molecule. **d**, The L510 dimer interface is stabilized by polar residues that are not part of the L510-LayV G interface. **e**, BLI measurements of mutants E24K and E27A. **f**, Prediction of the partial diffused MeV H binder M131pd5 superimposed with the prediction of the original M131 shows slightly altered binding pose. **g**, BLI measurement of M131pd5. **h**, ELISA of M131pd5 against a panel of paramyxovirus RBDs and unrelated controls. EC_50_ is reported for MeV H.

### L510 forms a homodimer that interferes with target binding

For protein characterization, we sought to experimentally determine the structures of the designs to further confirm their structural integrity. To our surprise, both elucidated *apo* binder structures, L510 and N237, were homodimers (Fig. 3c, Supplementary Fig.5b). The structure of N237 revealed a partially misfolded protein, forming a helix–turn–helix motif rather than the intended three-helix bundle. However, when paired with its neighboring molecule in the crystal, the resulting dimer still presents the designed binding interface, in principle enabling the predicted interaction without major steric clashes (Supplementary Fig. 5b). As N237 seemed to elute as a monomer from size exclusion chromatography, we expect this conformation to be a crystallographic artefact (Supplementary Fig. 4a). In contrast, L510 eluted with a shorter retention time, suggesting oligomerization. Indeed, the *apo* crystal structure confirmed the dimeric state of L510. The dimer interface consists of a hydrophobic patch and several hydrogen bonds formed by surface-exposed residues that were not part of the designed interface (Fig. 3d). Superimposed to the predicted complex, the second dimer molecule clashes into the target protein. Therefore, we hypothesized that the L510 *apo* dimer must dissociate prior to LayV G binding. To shift the equilibrium towards the LayV G bound state, we introduced mutations in the dimer interface to weaken the dimer formation. These allosteric mutations, E24K and E27A, exhibited up to 4-fold K_D_ improvement, highlighting the benefits of thorough characterization allowing for allosteric improvements (Fig. 3e).

### Partial diffusion can result in diverse sequences with improved affinity

Frequently seen in binder design campaigns is the sequence redesign of a functional scaffold using partial diffusion.^66,67^ Starting with a backbone ensemble generated by RFdiffusion, the possible sequence space increases and diverse interface sequences can be selected (Supplementary Fig. 6b). We applied this workflow to binders N237, L231, L510, and M131. For all scaffolds new functional sequences were identified, the success rates were similar to the initial *de novo* binder designs regarding specific and unspecific designs, and only one case resulted in an improved binder (Supplementary Fig. 7, 8). Design M131pd5, derived from M131, bound MeV H with a K_D_ of 36.5 nM in the BLI measurement, corresponding to a 6-fold improvement (Fig. 3g). The potency of MeV H detection in the ELISA increased as well, while also improving the specificity of the binder (Fig. 3h). Compared to the prediction of its parent, M131pd5 especially differed in its predicted loop region within the β-sheet, adopting an alternative conformation on the protein surface (Fig. 3f). Within the partial diffused mini proteins, N237pd4, which originated from our most promising binder candidate N237, exhibited the highest affinity with a K_D_ 32.6 nM (Supplementary Fig. 8).

### N237 competes with human ephrin B2 and neutralizing antibody for same epitope but only protects weakly against infectious NiV

We sought to further validate the functionality of our binders, focusing on the most potent candidate, N237, as our test case. To confirm the binding site as designed and predicted, we performed several competition assays. First, we evaluated the capability to inhibit human ephrin B2 binding to NiV G, as this was the intended design strategy (Fig. 4a). With increasing concentration of N237, the interaction between viral RBP and human receptor decreased (Fig. 4b), indicating direct competition, a strong surrogate of neutralizing activity. We further confirmed the binding site with two different antibodies. In contrast to HENV-32, which binds a different epitope, the neutralizing antibody HENV-117 competed with N237 for the receptor-binding site (Fig. 4c).^30^ Next, we sought to test whether these competitive properties survived stress tests, as the stability of *de novo* binders is often stated to be a major benefit in biotechnological applications.^42,46^ The competition against antibody HENV-117 was not affected by heat treatment for 15 min at 95°C, storage for 10 days at room temperature or even lyophilization without protection agent (Fig 4d). However, assessing the therapeutic potential, can only be evaluated by actual neutralization and protection of cells against infectious NiV.

**Fig. 4.:**
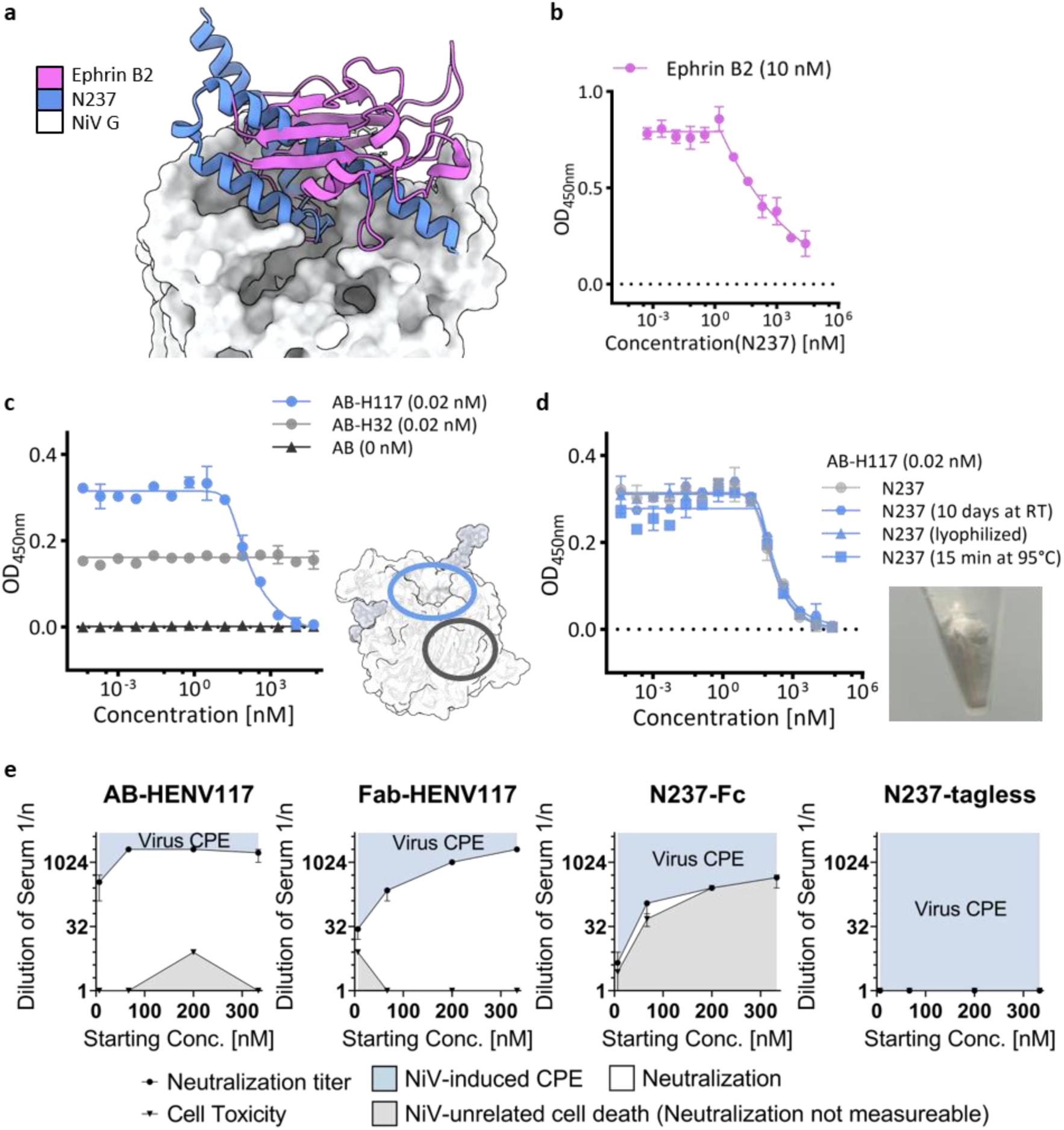
Validation of the epitope, stability and functionality of N237. **a**, Superimposed N237-NiV G prediction and ephrinB2-bound NiV G (PDB 2VSM) show overlapping binding sites on the RBD. **b**, Competition ELISA confirms the inhibition of receptor binding in the presence of increasing N237 concentration. **c**, Competition ELISA of N237 with antibodies HENV-117 (AB-H117) and HENV-32 (AB-H32). HENV-117, but not HENV-32, shows a decrease in response with increasing N237 concentration. **d**, Stress tested N237 binder (10 d at RT, lyophilized as shown in the image on the right, or 15 min at 95°C) retains function in HENV-117 competition. **e**, Neutralization assay of infectious NiV using Vero76 cells. The starting concentration of the dilution series plotted against titer of neutralization and titer of observed cell toxicity, both reported as the dilution step (n) of serum until the effect was observed. Error bars represent titer range in four technical replicates. The blue area shows NiV cytopathic effect (CPE), white area represents observed neutralization, grey area shows NiV unrelated cell death (Cell Toxicity = Neutralization not measurable).

Our neutralization assay with Fc tagged N237 suggests that despite the promising ELISA EC_50_ compared to HENV-117 (Fig. 2a, Supplementary Fig. 3f), the design cannot compete with the antibody regarding conclusive protection. (Fig. 4e). With a starting concentration of 66.7 nM, HENV-117 as antibody exhibits strong protection up to a dilution of 1:2048, corresponding to a concentration of 32.5 pM (Fig. 4e). In contrast, only weak neutralization by Fc-tagged N237 was detected at starting concentrations of 6.67 nM and 66.7 nM. Starting at 66.7 nM, dilutions of 1:8– 1:32 showed NiV-unrelated cell death, whereas dilutions >1:128 exhibited NiV-associated CPE.. Accordingly, N237 with a starting concentration of 66.7 nM displayed neutralization at dilutions of 1:64 (1.04 nM) and 1:128 (0.52 nM) (Fig. 4e). As our Fc tagged binder (N237-Fc and N101-Fc) induced cell toxicity at higher starting concentration, leading to inconclusive infection assessment, the neutralization titer within these dilution series cannot be measured (Fig. 4e, Supplementary Fig. 9). However, because N101 with an alternative Fc tag (Supplementary Table 1), identical to that used for HENV-117, as well as our tagless variants, did not show such toxicity, we conclude that the binders themselves do not induce cell death. The higher protective titer of Fc tagged HENV-117 and N237, compared to the Fab and the tagless variant, respectively, demonstrates the benefit of dimerization in virus neutralization. Conclusively, N237 displayed favorable properties for diagnostics or therapeutics, but ultimately failed in conclusive protection of viral infections compared to the control antibody.

### Predictive power of Boltz2 to distinguish binders and non-binders depends on the target protein

We retrospectively analyzed all our validated designs, excluding point mutants. All 67 designs comprising 19 binders, 5 unspecific binders, and 43 non-binders were predicted with AF2ig, our prediction tool used for majority of the designs, and Boltz2 combined with ipSAE calculation as state-of-the-art method for scoring (Supplementary Fig. 10).^68,69^ As expected, due to the filtering and design selection based on AF2ig LIS or pAE, LIS derived from AF2ig cannot distinguish between true positives and negatives within our data set. In contrast, Boltz2 confidences span a wider range and all 13 binders with a low ipSAE score (< 0.35) are non-binders. Interestingly, the discrimination of binders, unspecific binders, and non-binders via ipSAE was observed only for MeV H. Even the submicromolar binders pose slightly higher confidences than micromolar binders for MeV H (Supplementary Fig.10b). In the case of NiV G and LayV G, Boltz2 and AF2ig are both relatively confident in all binders, scoring non-binders equally well to binders, and are therefore unable to classify them.

## Discussion

We demonstrate that modern computational protein design tools, such as RFdiffusion and BindCraft, enable the *de novo* generation of mini-proteins targeting the RBP of paramyxoviruses. By combining *in silico* design with low-throughput *in vitro* screening, we establish a conceptual framework for a pandemic-preparedness platform. This platform integrates a computational module for candidate generation and filtering with an experimental module to validate binding to the RBDs of emerging viruses. 10–16 candidates were selected from the respective computational libraries per target protein, expressed recombinantly and characterized functionally. We identified specific binders against NiV G, MeV H, and LayV G, with affinities reaching into the low-nanomolar range. However, direct identification of high affine binders depended on the target protein and they varied largely in their biophysical properties (Supplementary Fig. 10a). Additionally, false positives were identified encouraging further work on improving methods and workflows to more rational and understandable *de novo* protein design. We further saw that different measurement types and fusion tags can influence properties, such as binding affinity. Most of the characterized hits exhibit exceptional protein expression yield, stability and specificity, demonstrating promising properties suitable for diagnostic tools. However, we did not confirm the binder’s potency in patient-derived samples, nor have we compared the minimal detection limit.

Additionally, we showed that even though N237 can compete for receptor binding with NiV G to human ephrin B2, the mini-protein only weakly neutralizes viral infections *in vitro* compared to the antibody HENV-117. The interaction of HeV G and NiV G with the human receptor ephrin B2 has been characterized as a strong interaction, especially when comparing the magnitude lower affinity of MeV H and SLAM.^51^ Potentially, even more affine competitive antibodies or binders for NiV are required than for other viruses. We hypothesize that higher affinities and multivalent display, to match the tetrameric RBP, are strategies to improve neutralization activity, as this has been used for other mini-proteins binding to viral glycoproteins such as MERS-CoV spike.^42^ In Ragotte et al., the authors demonstrate that trimeric constructs of the mini-protein cb3, improved the K_D_ value from 3.7 nM up to 21 pM against the trimeric glycoprotein. Additionally, all multivalent constructs exhibited higher neutralization than the monomeric mini-protein.

We also assessed computational affinity maturation strategies, ranging from point mutations to entirely new interfaces to allosteric mutations. We had moderate success with the introduction of single mutations to increase affinity. Further iterations would be required to improve K_D_ values. However, achieving low nanomolar or sub-nanomolar binding might require generation of new interface sequences with partial diffusion, screening of more *de novo* binders until a mini-protein with the desired affinity is identified. Alternatively, to the workflow presented here affinity maturation by laboratory directed evolution and display such as yeast display can be applied. Here, the selection pressure applied by cell sorting, in context of the binding response, will result in improved affinities, as shown by Ragotte et al.^42^ Compared to computational protein design where the majority of the prescreening is performed *in silico*, high-throughput display systems are time-consuming, labor intensive, costly and limited to laborites with such resources.

It might also be that our computationally designed protein backbones are too ideal and lack plasticity to accommodate a more diverse sequence profile, arguing that each scaffold would have a natural limitation for binding. It is likely that current research in generative models to design antibody-like protein binders will further increase the potential of computational design of highly potent protein-protein interactions.^70,71^ These upcoming state of the art tools can be incorporated as the *in silico* module of our platform, leading to a workflow with an even higher success rate.

Although we determined that our approach results in functional binders, we also demonstrated that *de novo* binder design requires thorough validation, including structure determination, to support the design hypothesis. The elucidated structure of L510 revealed an unwanted dimeric assembly of the mini-protein, proposing allosteric mutations in the dimer interface that improved the affinity 4-fold. Consequently, the experimental K_D_ value of the original L510 is biased by a property that was not part of the prediction in the design process, as only one mini-protein molecule is used in complex predictions.

In the context of application-oriented functional assessment, the *de novo* binders demonstrated high specificity and stability, making them well-suited for diagnostic tools such as lateral flow tests.^72^ We showed that N237’s competition against HENV-117 for binding to NiV G is unaffected by room-temperature storage for 10 days or lyophilization, supporting greater robustness of diagnostics under variable environmental conditions.^72,73^ Importantly, stability at ambient or high temperature not only addresses temperature challenges but also reduces the resources needed for cold-chain transport and storage, facilitating fairer and broader distribution of diagnostics.^74,75^

Similar advantages apply to therapeutic use. While N237 showed promising competition and neutralization activity, further optimization is needed. Testing of other NiV binders, affinity-matured N237_S47D, Fc-tagged variants, or multimeric constructs against infectious NiV in BSL-4 conditions is beyond the scope of this study, as we do not anticipate improvements beyond the antibody HENV-117. Nevertheless, our results confirm that mini-protein binders can neutralize viral infections. Combined with their diagnostic relevance across three paramyxovirus RBPs, this highlights the potential of such platforms to significantly enhance pandemic preparedness for an emerging disease X.

## Methods

### Computational design of *de novo* binders

Each target protein was analyzed individually to determine hotspot residues, mainly focusing on hydrophobic patches with adjacent polar residues, and forbidden interface residues, such as glycosylation sites. For each binder design campaign, up to 20 parameter configurations, including different input structures, input structure truncations, hotspot residues, RFdiffusion denoising configurations, ProteinMPNN configurations and the number of sequences per backbone, were evaluated by generating 100 binder backbones and subsequently subjected to sequence generation and complex prediction. Configurations that yielded the highest proportion of binders with high pLDDT and low pAE interaction were used for large scale *in silico* binder library generation. Using the RFdiffusion beta complex model, 2000 – 5000 backbones of 75-100 residues were diffused.^47^ Scaffolds were filtered prior to sequence generation. ProteinMPNN was used to generate 5-15 sequences per backbone at a temperature of 0.1.^39,76^ Alphafold2-initial guess (AF2ig) predictions with pAE interaction values < 15-20 were retained for sequence optimization.^39,77^ For each RFdiffusion backbone, a maximum of 2-5 binder sequences, ranked by pAE interaction, was retained. Sequence optimization was performed iteratively using ProteinMPNN on the predicted complex, increasing sampling to 15-20 sequences per backbone and temperature to 0.3. Iterations continued until no further pAE interaction improvement was observed or unitl only few backbones remained improving. The resulting *de novo* binder library was pre-filtered using AF2ig confidence metrics, pAE interaction or local interaction score (LIS).^63^ Accepted designs were further evaluated using the Rosetta Interface Analyzer (beta_nov16 scoring function) to calculate the interface energy (interface dG), shape complementarity, number of hydrogen bonds, solvent-accessible surface area, and determine interface residues and their interacting amino acid.^78^ Designs were selected by integrating multiple metrics and retaining binders within defined quantile thresholds (top 10% for pAE interaction and interface dG; top 25% for all other metrics). Selection criteria for each target included:

NiV G: AF2ig-pLDDT, AF2ig-pAE interaction, Rosetta-interface dG.

LayV G: AF2ig-pLDDT, AF2ig-LIS, Rosetta-interface dG, hydrogen bonds/ interface SASA, and frequency of recurrent interaction pairs for binders L63, L93, L94, L231, L389.

MeV H: AF2ig-pLDDT, AF2ig-LIS, Rosetta-interface dG, shape complementary, and frequency of recurrent interaction pairs in each binder.

Binder design using BindCraft was performed by first screening various preset advanced settings, target residues, and input structures to identify viable configurations.^38^ For LayV G binder design, BindCraft v1.1 was applied and resulted in no final accepted but several designs being rejected due to unsatisfied hbonds or surface hydrophobicity. From these designs, binders were selected by ipTM and visual inspection.^77^ For MeV H binder design, the same approach was applied using BindCraft v1.5, yielding 99 accepted design for the selection.

Visualization of predicted or experimentally determined protein structures was performed using ChimeraX.

### Computational optimization of binder hits

#### Partial diffusion - Full sequence redesign

Partial diffusion was applied to binders N237, L231, L510, M131. The AF2ig-predicted complex was used as input, in which the binder was first noised (partial_T = 15) and subsequently denoised (noise_scale = 0).^47^ A total of 500 backbones were generated, followed by the sequence design using ProteinMPNN to produce 25 sequences per backbone (temperature = 0.2) and complex prediction with AF2ig. Designs were scored as described for the initial *de novo* binder design workflow, with additional prioritization for sequence diversity within the selection, determined by multiple sequence alignment.

#### Consensus sequence – Point mutations via the frequency of sampled residues

Generated and filtered partially diffused binder libraries were analyzed individually on the consensus interface sequence for each binder. Positions with low diversity, where the most frequent amino acid was not the original amino acid, were taken as possible mutations. Four mutations for N237 (L4S/E7R, R11K, F44N, S47D) and one mutation for N101 (Y17Q) were selected.

#### Rosetta Mutational Scan – Point mutations via Rosetta interface dG

The predicted complex of N237 was energy minimized using a Rosetta FastRelax.^79^ Five relaxed protein complexes, selected randomly from the 20 best total scores out of 100 relaxations, were used further. In the mutational scan, each interface residue was mutated to all possible amino acids. For each input structure, five separate neighborhood relaxes were applied and the interface dG was calculated. N237 interface mutations with the largest improvement were selected for experimental validation (S47M, D54R, L58W, I65V).

#### LISxSC mutational scan – Point mutations via AF2ig-LIS and shape complementarity

ProteinMPNN was used to determine possible N237 interface mutations. Each possible point mutant was predicted five times using AF2ig, followed by analysis of the shape complementarity. The four mutations with the highest improvement of AF2ig-LIS and shape complementarity (E15D, D40R, L60I, K64Q) were selected for validation.

### LayV G head prediction using Colabfold

Crystal and CryoEM structures of LayV G (PDBs: 8K80, 8TVI, 8VWP) were used as templates to remodel the structure of the RBD using Colabfold.^25,61,77^ The model with the highest confidence (pTM) was used as binder design input.

### Prediction of binder-target complex using Boltz2

Each binder was predicted with its on-target using Boltz2.^80,81^ The structure of the targeted RBD was provided as template, the flag “use_potentials” was set true and no MSA was performed. For each predicted complex, ipSAE_min was calculated from the PAE matrix.^69^

### Gene synthesis and cloning

Genes, codon optimized for *Homo sapiens,* were ordered from Integrated DNA Technologies (IDT) as eBlocks™ with overhangs compatible with our in-house expression vector pCD_SEC2, which contains a tPA leader sequence. The pCD_SEC2 vector was modified to include a C-terminal GGGSG or GGGSGGGSG linker and human Fc tag. Genes were cloned using the NEBuilder HiFi DNA Assembly Cloning Kit (New England Biolabs). For expression in *E. coli*, genes were not further codon optimized and were amplified from the pCD_SEC2 vector using primers that contained overhangs for a modified pET11a vector with an N-terminal 6xHis tag and a TEV cleavage site. Genes and vector were assembled as described above.

All plasmids were validated by Sanger sequencing (Genewiz, Azenta).

### Recombinant protein expression

Viral glycoproteins and Fc tagged binders were expressed in Expi293 cells (ThermoFisher). Transfection was performed at a density of three million cells per milliliter using 1 µg DNA per mL culture, mixed with polyethlylenimine and Opti-MEM (ThermoFisher) in a ratio of one to three. Cells were incubated at 37°C, 8% CO_2_, humidified air and 200 rpm for tubes and flasks or 900 rpm for expression in 24-well plates. After 24 hours, 4 mg mL^-1^ glucose and 3 mM valproic acid were added and cells were incubated for an additional 3 to 6 days at either 33°C or 37°C, 8% CO_2_, humidified air and at 200 rpm or 900 rpm. Expressed proteins were harvested by centrifugation at 4000 rcf and 4°C for 20 min. Screening experiments and protein purifications were performed using the resulting supernatant.

For tagless binder production chemically competent *E. coli* BL21(DE3) cells (New England Biolabs) were used. Approximately 100 ng of plasmid containing the gene of interest were transformed into 25 µL of cells using heat shock. 10 mL LB, supplemented with 100 µg×mL^-1^ ampicillin, were inoculated and incubated overnight at 37°C and 220 rpm. This overnight culture was added to 800 mL LB-Amp and cells were grown to an OD_600_ of 0.6-0.8 before induction with 0.5 mM IPTG. Expression was proceeded at 19°C and 220 rpm for 14-18 hours. Cells were harvested by centrifugation at 9000*g* and 4°C for 20 min. Pellets were resuspended in immobilized ion metal chromatography (IMAC) buffer (50 mM HEPES pH 8.0, 300 mM NaCl) supplemented with DNase1 and lysozyme. Lysis was performed by sonication with 12 pulses of 10 s at 60% amplitude on ice. The lysate was cleared by centrifugation at 14000*g* and 4°C for 40 min. Protein purification was performed using the resulting supernatant.

### Protein purification

Fc tagged binders and antibodies were purified using a 1 mL MabSelect™ column (Cytiva) for affinity chromatography connected to a Masterflex Ismatec Peristaltic Pump (Avantor). The supernatant was loaded onto the column and washed with 8 mL tris-buffered saline (TBS) (20 mM TRIS pH 7.5, 150 mM NaCl). Proteins were eluted using 5 mL of 100 mM sodium citrate at pH 2.5, and the eluate was neutralized by adding 0.5 mL of 1 M TRIS at pH 8. The buffer was subsequently exchanged to TBS.

6xHis tagged proteins (viral glycoproteins and *E. coli* expressed binder) were purified using 0.5-2 mL Ni-NTA resin (Qiagen) for immobilized metal affinity chromatography. The supernatant containing the protein of interest was loaded onto the resin, which was washed twice with 15 mL of 50 mM HEPES pH 8.0, 1 M NaCl, and once with 20 mL IMAC buffer (50 mM HEPES pH 8.0, 300 mM NaCl). Elution was performed twice with 2.5 mL IMAC buffer supplemented with 400 mM imidazole. For binder purification, TEV cleavage was performed at a ratio of 1:100 (in-house purified TEV protease to protein of interest) at room temperature overnight. The eluate was applied to an ÄKTA pure purification system (Cytiva) for size exclusion chromatography (HiLoad 16/600 Superdex 75 pg (Cytiva) for binder, HiLoad 16/600 Superdex 200 pg (Cytiva) for glycoproteins) using 20 mM HEPES pH 8.0, 150 mM NaCl (HBS). Pure fractions were pooled and combined.

Molecular weight and purity of proteins were analyzed by SDS-PAGE, and protein concentration were determined by measuring the absorbance at 280 nm using a Nanodrop (ThermoFischer).

### Enzyme-linked immunosorbent assay (ELISA)

25 ng of viral glycoprotein, BSA, or DNA in 25 µL were coated per well in 384 high-binding plates using 100 mM NaHCO3 pH 9.6, followed by overnight incubation at 4°C. For dilution series ELISA to determine EC_50_ values, the amount of coated protein was increased to 250 ng for MeV H. After coating, wells were washed three times with TBS-T (TBS with 0.1% Tween 20) using a plate washer (BioTek EL406, Agilent). Blocking was performed by incubating wells with 50 µL TBS-T supplemented with 5% (w/v) milk powder for 60 min. After washing, binder and control antibodies (as purified protein or supernatant) were diluted in TBS-T supplemented with 2% (w/v) milk powder and incubated in the coated plates for 60 min. Wells were washed again. Peroxidase AffiniPure Goat Anti-Human IgG (H+L) (Jackson ImmunoResearch) was diluted 1:5000 in TBS-T supplemented with 2% (w/v) milk powder, and 25 µL were added per well and incubated for an additional 60 min. After three washes, binding was detected by adding 25 µL TMB reagent, and the reaction was quenched after 60-180 s by adding 25 µL of 1 M HCl. Absorbance was measured at 450 nm (BioTek Synergy H1, Agilent). Graphs are plotted using GraphPad Prism.

### Competition ELISA

Competition ELISA of binders with antibodies H117 and H32 against NiV G was performed using the same protocol as the standard ELISA.^82^ For the initial binder or antibody incubation, a dilution series of non-Fc tagged protein (tagless) was prepared in TBS-T supplemented with 2% (w/v) milk powder and mixed 1:1 with an Fc tagged antibody at a constant concentration. As a control, the Fc tagged protein at the same constant concentration was mixed 1:1 with buffer. Graphs are plotted using GraphPad Prism.

Human ephrin B2 receptor (hEFNB2) competition ELISA was performed analogous to the antibody competition. NiV G was coated at a concentration of 10 ng µL^-1^. The ephrin B2 concentration was constant at 10 nM and N237 tagless was diluted starting at 25 µM. Ephrin B2 binding was observed by detection of its His tag using an anti-his HRP conjugate (QIAGEN). Graphs are plotted using GraphPad Prism.

### Biolayer interferometry (BLI)

Purified FC tagged proteins were diluted to 2 µg mL^-1^ in 10 mM HEPES pH 7.4, 150 mM NaCl, 3 mM EDTA, 0.1% BSA, 0.05% Tween 20. For screening experiments, Expi293 supernatant was diluted 1:50, and for K_D_ measurements using unpurified protein the dilution was adjusted to achieve loading levels comparable to the purified protein. BLI experiments were performed on an Octet R8 (Sartorius) using AHC or Protein A biosensors (Sartorius). Diluted binder or antibody samples were loaded on the biosensor for 90 s, followed by a 60 s baseline. Association, either at a single concentration for screening or a dilution series of viral glycoprotein for kinetics, was recorded for 300 s. Dissociation was performed for 300 s. Biosensors were regenerated in 0.85% phosphoric acid up to 12 times. BLI Data analysis was performed using the provided analysis software. The reference well containing no analyte in the association step was subtracted and data were aligned to 55.0-59.8 s of the baseline. Inter-step correction to dissociation and Savitzky-Golay filtering were applied. Data were fitted globally with a 1:1 binding model. If the fit was low quality (Full_R^2^ < 0.99), either the dissociation time was reduced to improve the fit of the initial dissociation phase (> 10% of dissociation) or steady state analysis was applied.

### Circular dichroism (CD)

Proteins were diluted in 5 mM KPi pH 7.4, 25 mM Na_2_SO_4_ to a concentration of 20 µg mL^-1^. A J-1500 CD spectrometer (Jasco) was used to measure the ellipticity at 200-260 nm across temperature intervals. Starting at 25°C, full spectra were recorded at 5°C intervals up to 95°C with a temperature ramp of 1°C min^-1^ applied.

### Analytical Size Exclusion Chromatography

Approximately 1-2 mg of protein in 500 µL HBS were injected onto a SD75 Increase 10/300 GL (Cytiva) for binders and a SD200 Increase 10/300 GL (Cytiva) for the viral glycoproteins using an ÄKTA pure purification system.

### Isothermal titration calorimethry (ITC)

Proteins, polished via analytical SEC into the identical HBS, were used for ITC measurements (MicroCal PEAQ, Malvern). Viral glycoproteins were diluted to 10 µM and binders to a concentration of 100 µM. The sample cell was loaded with viral glycoprotein. Within 12 or 19 injections, each separated by 150 s spacing, the binder was titrated into the cell up to a molar ratio of two. Temperature during the experiment was set at 25°C, reference power to 5 µcal s^-1^, Feedback to high and stir speed to 750 rpm. The ITC Analysis Software (v1.41, Malvern) was used for peak integration and baseline subtraction.

### Protein lyophilization

9 mg of N237, without tag, in 500 µL HBS were lyophilized using an Alpha 2-4 LSCbasic (Christ) freeze-dryer. The powder was stored at -20 °C for 10 d before resuspension with water to 500 µL.

### Protein crystallization, data collection and structure determination

Protein crystals of N237 *apo* were obtained at a concentration of 8 mg mL^-1^ using hanging drop vapor diffusion at 24°C in 30% (w/v) PEG3350 and 0.2 M NaCl. The protein and reservoir solution were mixed in a one-to-one ratio. L510 *apo* was crystallized using the hanging drop vapor diffusion method by combining 1 µL protein in HBS at a concentration of 20 mg mL^-1^ and 1 µL reservoir solution (60% (w/v) MPD, 0.1 M HEPES pH 7.5). Crystals were cryoprotected using 25 % (v/v) ethylene glycol before flash-freezing in liquid nitrogen.

Crystallographic data were collected at BESSY II Beamline MX14.1 and EMBL/DESY Beamline P14, respectively.^83^ Diffraction data were processed using the autoPROC package and phases were determined by molecular replacement using the predicted binder structure as search model.^84–86^ Model building was performed manually in Coot, and automated refinement was carried out with PHENIX.refine.^87,88^ Data collection, phasing, and refinement statistics are summarized in Table S1.

### Cells and virus

Vero76 cells (Collection of Cell Lines in Veterinary Medicine, Friedrich-Loeffler-Institut, FLI; CCLV-Rie 0228) were maintained in Dulbecco’s modified Eagle’s medium (DMEM) supplemented with 10% fetal calf serum and incubated at 37°C.

NiV (NiV Malaysia, GenBank accession no AF212302) was propagated and titrated on Vero76 cells by plaque assay as described previously with slight modifications.^89^ Work with live NiV was performed at the BSL4-facility of the Friedrich-Loeffler-Institut, Greifswald – Insel Riems.

### Determination of cell toxicity

To measure cell toxicity in Vero76 cells, an LDH-Assay (Promega) was performed according to the manufacturer’s protocol. Briefly, white microtiter plates were equipped with LDH-storage buffer and the supernatants of treated cells were added in a 1:25 ratio. Plates then were either stored at -20 °C until measuring or immediately measured. For this, LDH Detection Reagent was prepared by combining the LDH Detection Enzyme Mix and Reductase Substrate and this mix was added to each well containing the LDH-storage buffer and the supernatants. The reaction was allowed to proceed for 1 h at room temperature in the dark before luminescence was measured using a luminometer (TriStar). For calculating cell toxicity, cells treated with 1%Triton served as positive control and these values were set as 100%.

### NiV neutralization assay

To measure the neutralizing activity of the various mini-proteins variants and control, HENV-117 as antibody and Fab, the starting concentrations of 6.67 nM, 66.7 nM, 200 nM, and 300 nM were two-fold serially diluted in a 96-well plate. Then, 25 plaque forming units (PFU) of NiV were added to each well. After incubation at 37°C for 1 h, 9x10^3^ Vero76 cells were added to each well and plates were incubated further at 37°C for 4 days. Then, CPE was visually scored and neutralizing titer determined. Each dilution series, with different starting concentration, was performed with four replicates (4x 4 dilution series) and titers were averaged. Graphs are plotted using GraphPad Prism.

## Data and Software Availability

Rosetta is freely available for non-commercial use (www.rosettacommons.org). All structural biology software used for molecular replacement and refinement are part of SBGrid.^90^ Supplementary material, including experimental data, computational scripts, and all tested sequences, will be made publicly available with the peer-reviewed publication.

## Supplementary Information

**Supplementary Fig. 1:** Structure of NiV, LayV and MeV RBP.

**Supplementary Fig. 2:** Screening of *de novo* designed binders.

**Supplementary Fig. 3:** Binding experiments of positive hits, control, and unspecific binder.

**Supplementary Fig. 4:** Supplementary characterization of selected NiV, LayV and MeV RBP binders.

**Supplementary Fig. 5:** Specificity of NiV G binders and crystal structure of N237.

**Supplementary Fig. 6:** Strategies for the *in silico* optimization of binder hits.

**Supplementary Fig. 7:** Screening of partial diffused binders.

**Supplementary Fig. 8:** Affinity determination of partial diffused binder and N237 point mutants.

**Supplementary Fig. 9:** Neutralization and cell toxicity assay

**Supplementary Fig. 10:** Retroperspective AF2ig and Boltz2 analysis of experimentally characterized binder.

**Supplementary Table 1.** Sequences of the characterized binders.

**Supplementary Table 2.** Sequences of glycoproteins, antibodies and ephrin B2. **Supplementary Table 3.** Crystallographic table.

## Contact for resource sharing

Further information and requests for computational and material resources and reagents should be directed to and will be fulfilled by the Lead Contact.

## Acknowledgements

We thank Sascha Wagner, Birke Brumme and Christina M. Lange for excellent technical assistance. We appreciate Moritz Ertelt for great discussions on computational protein design. The Leipzig University Computing Center provided computational resources. The synchrotron data was partly collected at beamline operated by EMBL Hamburg at the PETRA III storage ring (DESY, Hamburg, Germany). We also thank Dr. Renato Weiße and Prof. Norbert Sträter for their help regarding macromolecular crystallography, and Gert Weber and Manfred Weiss for their help at BESSY II, Helmholtz Zentrum Berlin. C.T.S further likes to thank Prof. Dr. Christine Engeland for spreading the excitement for Measles virus research.

## Author information

Authors and Affiliations

**Institute for Drug Discovery, Medical Faculty, Leipzig University, Leipzig, Germany**

Dominic Rieger, Antonia Sophia Peter, Phillip Schlegel, Anna Nobis, Hannes Junker, Lena Kiesewetter, Max Beining, Lorenz Beckmann, Johannes Klier, Jens Meiler, Clara T. Schoeder

**Friedrich-Loeffler-Institut, Institute of Novel and Emerging Infectious Diseases, Greifswald-Insel Riems, Germany**

Carolin Rüdiger, Sandra Diederich

**Division of Structural Biology, Centre for Human Genetics, University of Oxford, Oxford, United Kingdom**

Antonia Sophia Peter, Nichakorn Pipatpadungsin, Robert Stass, Thomas A. Bowden

**School of Embedded Composite Artificial Intelligence (SECAI), Cooperation of University Leipzig and TU Dresden**

Max Beining, Johannes Klier, Jens Meiler

**Center for Scalable Data Analytics and Artificial Intelligence ScaDS.AI, Dresden/Leipzig, Germany**

Jens Meiler, Clara T. Schoeder

**Department of Chemistry, Department of Pharmacology, Center for Structural Biology, Institute of Chemical Biology, Center for Applied Artificial Intelligence in Protein Dynamics, Vanderbilt University, Nashville, Tennessee, United States of America**

Jens Meiler

## Contributions

C.T.S and D.R. conceptualized the project and designed experiments. D.R., H.J., M.B., J.K developed the modified binder design pipeline. D.R. performed computational design and optimization of binders. D.R. A.N., L.K purified viral glycoproteins and performed screening experiments. D.R., P.S. characterized binders structurally and functionally. L.B., D.R. performed binder lyophilization. A.S.P, N.P., R.S. and T.A.B. designed and performed ephrin B2 competition assay. C.R. and S.D. designed and conducted Nipah virus neutralization assay. D.R, C.T.S and S.D. wrote the original draft of the manuscript. All authors read and finalized the manuscript and confirmed their contributions.

## References

1. Gahr, P. et al. An Outbreak of Measles in an Undervaccinated Community. Pediatrics 134, e220–e228 (2014).

2. Bankamp, B., Hickman, C., Icenogle, J. P. & Rota, P. A. Successes and challenges for preventing measles, mumps and rubella by vaccination. Curr. Opin. Virol. 34, 110–116 (2019).

3. Yang, L., Grenfell, B. T. & Mina, M. J. Waning immunity and re-emergence of measles and mumps in the vaccine era. Curr. Opin. Virol. 40, 48–54 (2020).

4. Chua, K. B. Nipah virus outbreak in Malaysia. J. Clin. Virol. 26, 265–275 (2003).

5. Chua, K. B. et al. Nipah Virus: A Recently Emergent Deadly Paramyxovirus. Science 288, 1432–1435 (2000).

6. Wu, Z. et al. Novel Henipa-like Virus, Mojiang Paramyxovirus, in Rats, China, 2012. Emerg. Infect. Dis. 20, (2014).

7. Mahalingam, S. et al. Hendra virus: an emerging paramyxovirus in Australia. Lancet Infect. Dis. 12, 799–807 (2012).

8. Chakraborty, S. et al. Langya virus, a newly identified Henipavirus in China - Zoonotic pathogen causing febrile illness in humans, and its health concerns: Current knowledge and counteracting strategies – Correspondence. Int. J. Surg. 105, 106882 (2022).

9. Shafaati, M. & Zandi, M. Langya henipavirus (LayV) as an emerging zoonotic disease: a mini-review. New Microbes New Infect. 68, 101643 (2025).

10. Caruso, S. & Edwards, S. J. Recently Emerged Novel Henipa-like Viruses: Shining a Spotlight on the Shrew. Viruses 15, 2407 (2023).

11. Luby, S. P. et al. Recurrent Zoonotic Transmission of Nipah Virus into Humans, Bangladesh, 2001–2007. Emerg. Infect. Dis. 15, 1229–1235 (2009).

12. Li, H., Kim, J.-Y. V. & Pickering, B. S. Henipavirus zoonosis: outbreaks, animal hosts and potential new emergence. Front. Microbiol. 14, 1167085 (2023).

13. Gazal, S. et al. Nipah and Hendra Viruses: Deadly Zoonotic Paramyxoviruses with the Potential to Cause the Next Pandemic. Pathogens 11, 1419 (2022).

14. Duprex, W. P. & Dutch, R. E. Paramyxoviruses: Pathogenesis, Vaccines, Antivirals, and Prototypes for Pandemic Preparedness. J. Infect. Dis. 228, S390–S397 (2023).

15. Thakur, N. & Bailey, D. Advances in diagnostics, vaccines and therapeutics for Nipah virus. Microbes Infect. 21, 278–286 (2019).

16. Gurley, E. S. & Plowright, R. K. A Roadmap of Primary Pandemic Prevention Through Spillover Investigation. Emerg. Infect. Dis. 31, (2025).

17. Gómez Román, R., et al. Nipah@20: Lessons Learned from Another Virus with Pandemic Potential. mSphere 5, 10.1128/msphere.00602-20 (2020).

18. Chan, X. H. S. et al. Therapeutics for Nipah virus disease: a systematic review to support prioritisation of drug candidates for clinical trials. Lancet Microbe 6, 101002 (2025).

19. Cox, R. M. & Plemper, R. K. Structure and organization of paramyxovirus particles. Curr. Opin. Virol. 24, 105–114 (2017).

20. Marcink, T. C. et al. Subnanometer structure of an enveloped virus fusion complex on viral surface reveals new entry mechanisms. Sci. Adv. 9, eade2727 (2023).

21. Lee, B. & Ataman, Z. A. Modes of paramyxovirus fusion: a Henipavirus perspective. Trends Microbiol. 19, 389–399 (2011).

22. Stelfox, A. J. et al. Molecular basis for occlusion of the jeilongvirus receptor-binding site by the elongated C-terminus. mBio e01501–25 (2025) doi:10.1128/mbio.01501-25.

23. Bowden, T. A., Crispin, M., Jones, E. Y. & Stuart, D. I. Shared paramyxoviral glycoprotein architecture is adapted for diverse attachment strategies. Biochem. Soc. Trans. 38, 1349– 1355 (2010).

24. Xu, K., et al. Host cell recognition by the henipaviruses: Crystal structures of the Nipah G attachment glycoprotein and its complex with ephrin-B3. Proc. Natl. Acad. Sci. 105, 9953– 9958 (2008).

25. Wang, Z. et al. Structure and design of Langya virus glycoprotein antigens. Proc. Natl. Acad. Sci. 121, e2314990121 (2024).

26. Wang, Z. et al. Architecture and antigenicity of the Nipah virus attachment glycoprotein. Science 375, 1373–1378 (2022).

27. Aguilar, H. C. et al. A Novel Receptor-induced Activation Site in the Nipah Virus Attachment Glycoprotein (G) Involved in Triggering the Fusion Glycoprotein (F). J. Biol. Chem. 284, 1628–1635 (2009).

28. Navaratnarajah, C. K. et al. The Measles Virus Hemagglutinin Stalk: Structures and Functions of the Central Fusion Activation and Membrane-Proximal Segments. J. Virol. 88, 6158–6167 (2014).

29. Iankov, I. D. et al. Neutralization capacity of measles virus H protein specific IgG determines the balance between antibody-enhanced infectivity and protection in microglial cells. Virus Res. 172, 15–23 (2013).

30. Doyle, M. P. et al. Cooperativity mediated by rationally selected combinations of human monoclonal antibodies targeting the henipavirus receptor binding protein. Cell Rep. 36, 109628 (2021).

31. Gao, Z. et al. Assessment of the immunogenicity and protection of a Nipah virus soluble G vaccine candidate in mice and pigs. Front. Microbiol. 13, 1031523 (2022).

32. Julik, E. & Reyes-del Valle, J. Generation of a More Immunogenic Measles Vaccine by Increasing Its Hemagglutinin Expression. J. Virol. 90, 5270–5279 (2016).

33. Zhu, W., Smith, G., Pickering, B., Banadyga, L. & Yang, M. Enzyme-Linked Immunosorbent Assay Using Henipavirus-Receptor EphrinB2 and Monoclonal Antibodies for Detecting Nipah and Hendra Viruses. Viruses 16, 794 (2024).

34. Salleh, M. Z. Structural biology of Nipah virus G and F glycoproteins: Insights into therapeutic and vaccine development. Eur. J. Microbiol. Immunol. 15, 83–93 (2025).

35. Sun, X. et al. A Highly Specific Antibody-Based Assay for Nipah Virus AlphaLISA Detection. Viruses 17, 748 (2025).

36. Li, W. et al. Structure and function of a pair of non-competing monoclonal antibodies against Langya henipavirus attachment glycoprotein. Cell Rep. 44, 116407 (2025).

37. Han, K. H., Li, Y.-C., Parveen, R., Venkataraman, S. & Lin, C.-W. Technologies for Monoclonal Antibody Discovery and Development. Int. J. Mol. Sci. 26, 10470 (2025).

38. Pacesa, M. et al. One-shot design of functional protein binders with BindCraft. Nature 646, 483–492 (2025).

39. Bennett, N. R. et al. Improving de novo protein binder design with deep learning. Nat. Commun. 14, 2625 (2023).

40. Cheng, J., Liang, T., Xie, X.-Q., Feng, Z. & Meng, L. A new era of antibody discovery: an in-depth review of AI-driven approaches. Drug Discov. Today 29, 103984 (2024).

41. Zheng, J., Wang, Y., Liang, Q., Cui, L. & Wang, L. The Application of Machine Learning on Antibody Discovery and Optimization. Molecules 29, 5923 (2024).

42. Ragotte, R. J. et al. Designed miniproteins potently inhibit and protect against MERS-CoV. Cell Rep. 44, 115760 (2025).

43. Chazin-Gray, A. M. et al. De Novo Design of Miniprotein Inhibitors of Bacterial Adhesins. Preprint at 10.1101/2025.08.18.670751 (2025).

44. Liu, B. et al. Design of high-specificity binders for peptide–MHC-I complexes. Science 389, 386–391 (2025).

45. Koga, R. et al. Robust folding of a de novo designed ideal protein even with most of the core mutated to valine. Proc. Natl. Acad. Sci. 117, 31149–31156 (2020).

46. Vázquez Torres, S., et al. De novo designed proteins neutralize lethal snake venom toxins. Nature 639, 225–231 (2025).

47. Watson, J. L. et al. De novo design of protein structure and function with RFdiffusion. Nature 620, 1089–1100 (2023).

48. Aguilar, H. C., Henderson, B. A., Zamora, J. L. & Johnston, G. P. Paramyxovirus Glycoproteins and the Membrane Fusion Process. Curr. Clin. Microbiol. Rep. 3, 142–154 (2016).

49. Bowden, T. A., Crispin, M., Harvey, D. J., Jones, E. Y. & Stuart, D. I. Dimeric architecture of the Hendra virus attachment glycoprotein: evidence for a conserved mode of assembly. J. Virol. 84, 6208–6217 (2010).

50. Rissanen, I. et al. Idiosyncratic Mòjiāng virus attachment glycoprotein directs a host-cell entry pathway distinct from genetically related henipaviruses. Nat. Commun. 8, 16060 (2017).

51. Bowden, T. A. et al. Structural basis of Nipah and Hendra virus attachment to their cell-surface receptor ephrin-B2. Nat. Struct. Mol. Biol. 15, 567–572 (2008).

52. Zhang, X. et al. Structure of measles virus hemagglutinin bound to its epithelial receptor nectin-4. Nat. Struct. Mol. Biol. 20, 67–72 (2013).

53. Hashiguchi, T. et al. Crystal structure of measles virus hemagglutinin provides insight into effective vaccines. Proc. Natl. Acad. Sci. 104, 19535–19540 (2007).

54. Bergeron, É. et al. Streamlined detection of Nipah virus antibodies using a split NanoLuc biosensor. Emerg. Microbes Infect. 13, 2398640 (2024).

55. Bowden, T. A., Crispin, M., Jones, E. Y. & Stuart, D. I. Shared paramyxoviral glycoprotein architecture is adapted for diverse attachment strategies. Biochem. Soc. Trans. 38, 1349– 1355 (2010).

56. Zeltina, A., Bowden, T. A. & Lee, B. Emerging Paramyxoviruses: Receptor Tropism and Zoonotic Potential. PLoS Pathog. 12, e1005390 (2016).

57. Hashiguchi, T. et al. Crystal structure of measles virus hemagglutinin provides insight into effective vaccines. Proc. Natl. Acad. Sci. 104, 19535–19540 (2007).

58. Hashiguchi, T. et al. Structure of the measles virus hemagglutinin bound to its cellular receptor SLAM. Nat. Struct. Mol. Biol. 18, 135–141 (2011).

59. Sappington, I. et al. Improved protein binder design using beta-pairing targeted RFdiffusion. Preprint at 10.1101/2024.10.11.617496 (2024).

60. Wang, C. et al. Structural insights into the Langya virus attachment glycoprotein. Structure 32, 1090–1098.e3 (2024).

61. Mirdita, M. et al. ColabFold: making protein folding accessible to all. Nat. Methods 19, 679– 682 (2022).

62. Abramson, J. et al. Accurate structure prediction of biomolecular interactions with AlphaFold 3. Nature 630, 493–500 (2024).

63. Kim, A.-R. et al. Enhanced Protein-Protein Interaction Discovery via AlphaFold-Multimer. Preprint at 10.1101/2024.02.19.580970 (2024).

64. Zhu, Z. et al. Exceptionally Potent Cross-Reactive Neutralization of Nipah and Hendra Viruses by a Human Monoclonal Antibody. J. Infect. Dis. 197, 846–853 (2008).

65. Dong, J. et al. Potent Henipavirus Neutralization by Antibodies Recognizing Diverse Sites on Hendra and Nipah Virus Receptor Binding Protein. Cell 183, 1536–1550.e17 (2020).

66. Vázquez Torres, S., et al. De novo design of high-affinity binders of bioactive helical peptides. Nature 626, 435–442 (2024).

67. Liu, C. et al. Diffusing protein binders to intrinsically disordered proteins. Preprint at 10.1101/2024.07.16.603789 (2024).

68. Overath, M. D. et al. Predicting Experimental Success in De Novo Binder Design: A Meta-Analysis of 3,766 Experimentally Characterised Binders. Preprint at 10.1101/2025.08.14.670059 (2025).

69. Dunbrack, R. L. Rēs ipSAE loquunt : What’s wrong with AlphaFold’s ipTM score and how to fix it. Preprint at 10.1101/2025.02.10.637595 (2025).

70. Mille-Fragoso, L. S. et al. Efficient generation of epitope-targeted de novo antibodies with Germinal. Preprint at 10.1101/2025.09.19.677421 (2025).

71. Bennett, N. R. et al. Atomically accurate de novo design of antibodies with RFdiffusion. Nature https://doi.org/10.1038/s41586-025-09721-5 (2025) doi:10.1038/s41586-025-09721-5.

72. Goyal, G., Sharma, A., Tok, A. I. Y., Palaniappan, A. & Liedberg, B. Affimer sandwich probes for stable and robust lateral flow assaying. Anal. Bioanal. Chem. 414, 4245–4254 (2022).

73. Ramachandran, S., Fu, E., Lutz, B. & Yager, P. Long-term dry storage of an enzyme-based reagent system for ELISA in point-of-care devices. The Analyst 139, 1456–1462 (2014).

74. Sharma, S., Zapatero-Rodríguez, J., Estrela, P. & O’Kennedy, R. Point-of-Care Diagnostics in Low Resource Settings: Present Status and Future Role of Microfluidics. Biosensors 5, 577–601 (2015).

75. Ganchi, F. A. & Hardcastle, T. C. Role of Point-of-Care Diagnostics in Lower- and Middle-Income Countries and Austere Environments. Diagn. Basel Switz. 13, 1941 (2023).

76. Dauparas, J. et al. Robust deep learning–based protein sequence design using ProteinMPNN. Science 378, 49–56 (2022).

77. Jumper, J. et al. Highly accurate protein structure prediction with AlphaFold. Nature 596, 583–589 (2021).

78. Stranges, P. B. & Kuhlman, B. A comparison of successful and failed protein interface designs highlights the challenges of designing buried hydrogen bonds. Protein Sci. 22, 74–82 (2013).

79. Conway, P., Tyka, M. D., DiMaio, F., Konerding, D. E. & Baker, D. Relaxation of backbone bond geometry improves protein energy landscape modeling. Protein Sci. 23, 47–55 (2014).

80. Wohlwend, J. et al. Boltz-1 Democratizing Biomolecular Interaction Modeling. Preprint at 10.1101/2024.11.19.624167 (2024).

81. Passaro, S. et al. Boltz-2: Towards Accurate and Efficient Binding Affinity Prediction. Preprint at 10.1101/2025.06.14.659707 (2025).

82. Peter, A. S., et al. Characterization of SARS-CoV-2 Escape Mutants to a Pair of Neutralizing Antibodies Targeting the RBD and the NTD. Int. J. Mol. Sci. 23, 8177 (2022).

83. Mueller, U. et al. The macromolecular crystallography beamlines at BESSY II of the Helmholtz-Zentrum Berlin: Current status and perspectives. Eur. Phys. J. Plus 130, 141 (2015).

84. Vonrhein, C. et al. Data processing and analysis with the *autoPROC* toolbox. Acta Crystallogr. D Biol. Crystallogr. 67, 293–302 (2011).

85. Kabsch, W. XDS. Acta Crystallogr. D Biol. Crystallogr. 66, 125–132 (2010).

86. Evans, P. Scaling and assessment of data quality. Acta Crystallogr. D Biol. Crystallogr. 62, 72–82 (2006).

87. Afonine, P. V. et al. Towards automated crystallographic structure refinement with *phenix.refine*. Acta Crystallogr. D Biol. Crystallogr. 68, 352–367 (2012).

88. Emsley, P., Lohkamp, B., Scott, W. G. & Cowtan, K. Features and development of *Coot*. Acta Crystallogr. D Biol. Crystallogr. 66, 486–501 (2010).

89. Weingartl, H. M. et al. Recombinant Nipah Virus Vaccines Protect Pigs against Challenge. J. Virol. 80, 7929–7938 (2006).

90. Morin, A. et al. Collaboration gets the most out of software. eLife 2, e01456 (2013).

