## Supplementary Information for "Computational design and experimental characterization of mini-protein binders targeting Nipah, Langya and Measles virus receptor-binding domains"

D. Rieger *et al.*

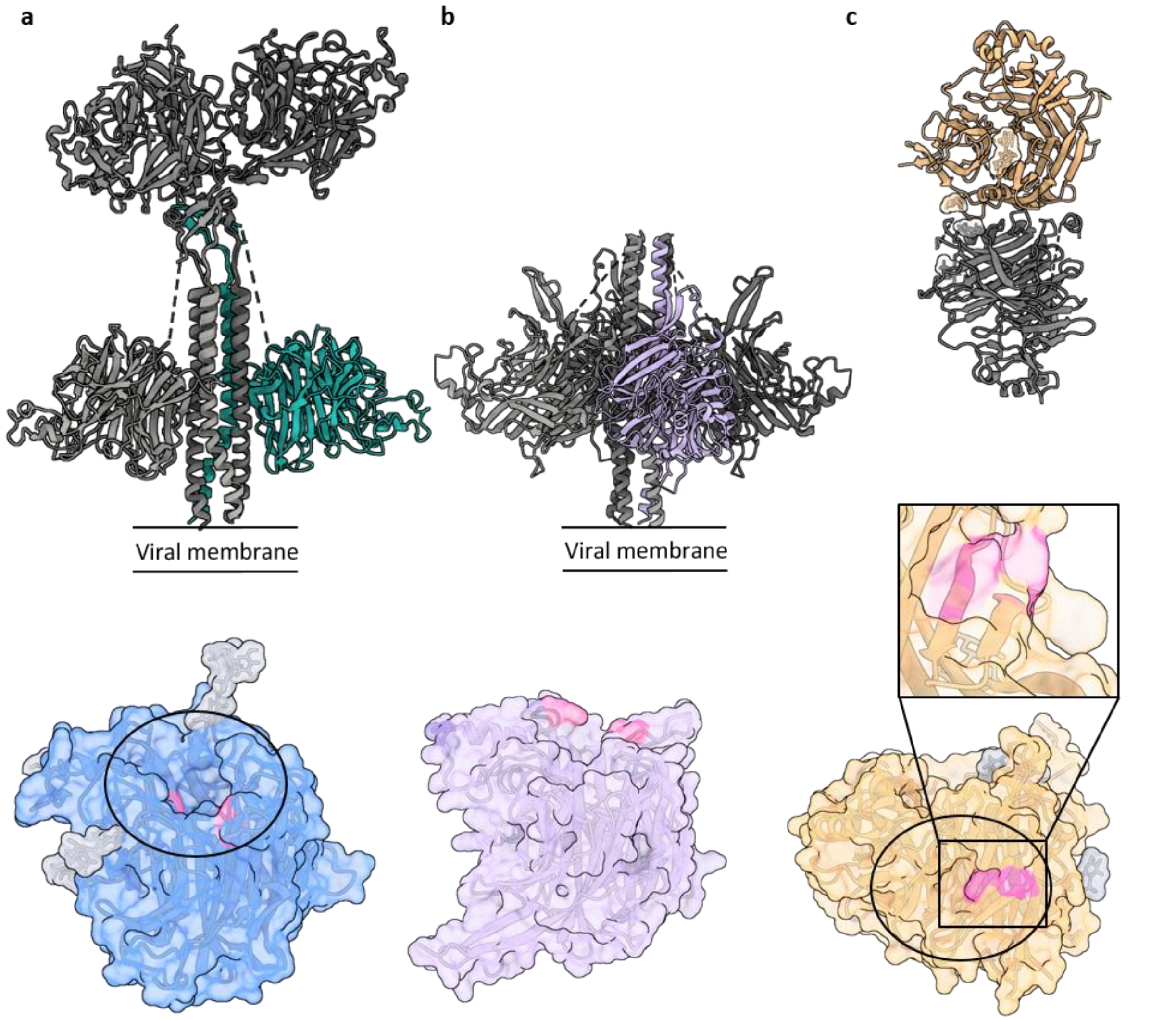


**Supplementary Fig. 1: Structure of NiV, LayV and MeV RBP.** The tetrameric structures of LayV G and NiV G display one possible conformation of a dynamic protein. Glycans are only shown on the RBD (bottom) in grey. **a**, Top: NiV G (PDB 7TY0, 7TXZ - superimposed) forms a homotetramer with two RBDs at the tip of the stalk and two RBDs adjacent to the stalk. Bottom: RBD of NiV G (PDB 3D11) with target residues marked in pink (F358, Y518, I588). The receptor-binding site is indicated by the black ellipse. **b**, Top: LayV G (PDB 8VWP) forms a homotetramer with four RBDs adjacent to the stalk. Bottom: Colabfold prediction of the LayV G RBD with target residues marked in pink (P311, M313, F403). **c**, Top: Crystal structure of dimerized MeV H RBD

(PDB 2ZB6) without its stalk. Bottom: RBD of MeV H with target residues marked in pink (Y541, F522). The receptor-binding site is indicated by the black ellipse.

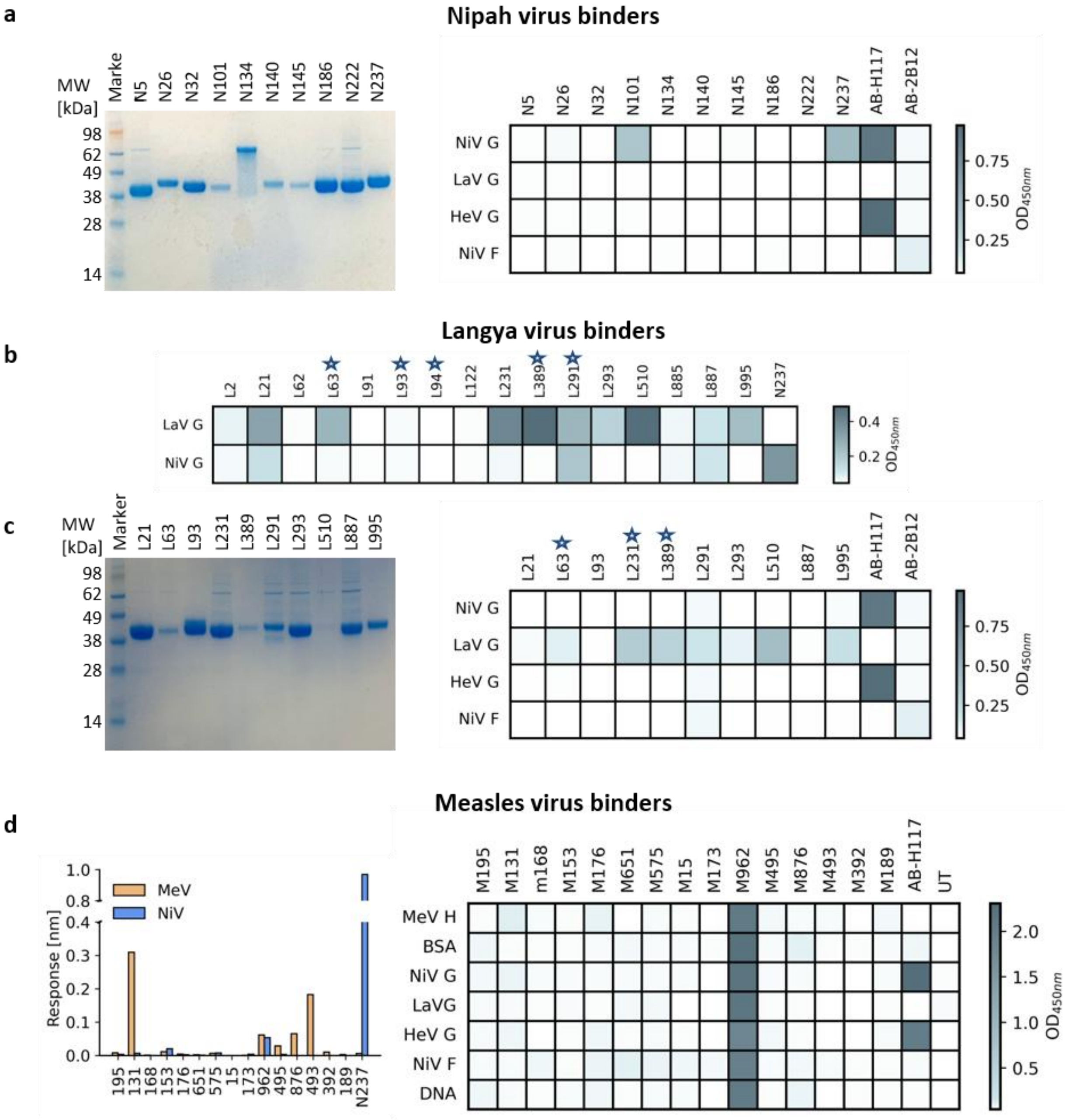


**Supplementary Fig. 2: Screening of *de novo* designed binders.** **a**, SDS-PAGE of purified Fc tagged NiV G binders and ELISA using 300 nM binder. **b**, ELISA of LayV G binder using unpurified protein. Binders, including frequency of interaction pairs as score, are marked with a star. **c**, SDS-PAGE of purified Fc tagged LayV G binders and ELISA using 300 nM binder. Binders, including frequency of interaction pairs as score, are marked with a star. **d**, BLI using immobilized Fc tagged MeV H binders and 1000 nM MeV H or NiV G for association (left). ELISA using 300 nM binder (right).

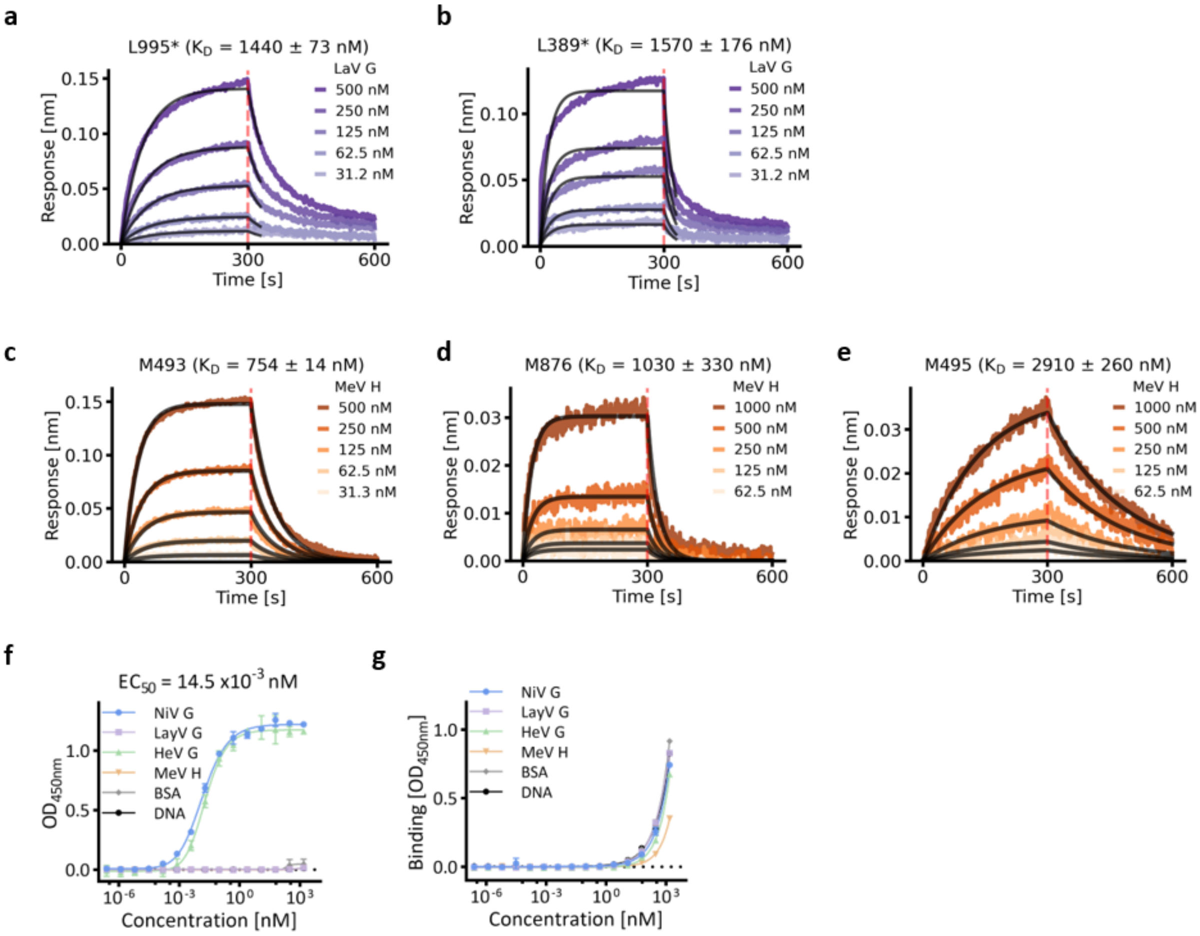


**Supplementary Fig. 3: Binding experiments of positive hits, control, and unspecific binder.** * indicates reduction of the dissociation time in post-processing to improve accuracy of the fit. **a**, BLI measurement of LayV G binder L995. **b**, BLI measurement of LayV G binder L389. **c**, BLI measurement of MeV H binder M493. **c**, BLI measurement of MeV H binder M876. **c**, BLI measurement of MeV H binder M495. **f**, ELISA of antibody HENV-117 against a panel of paramyxovirus RBDs and unrelated controls. **g**, ELISA of unspecific binder M962 against a panel of paramyxovirus RBDs and unrelated controls.

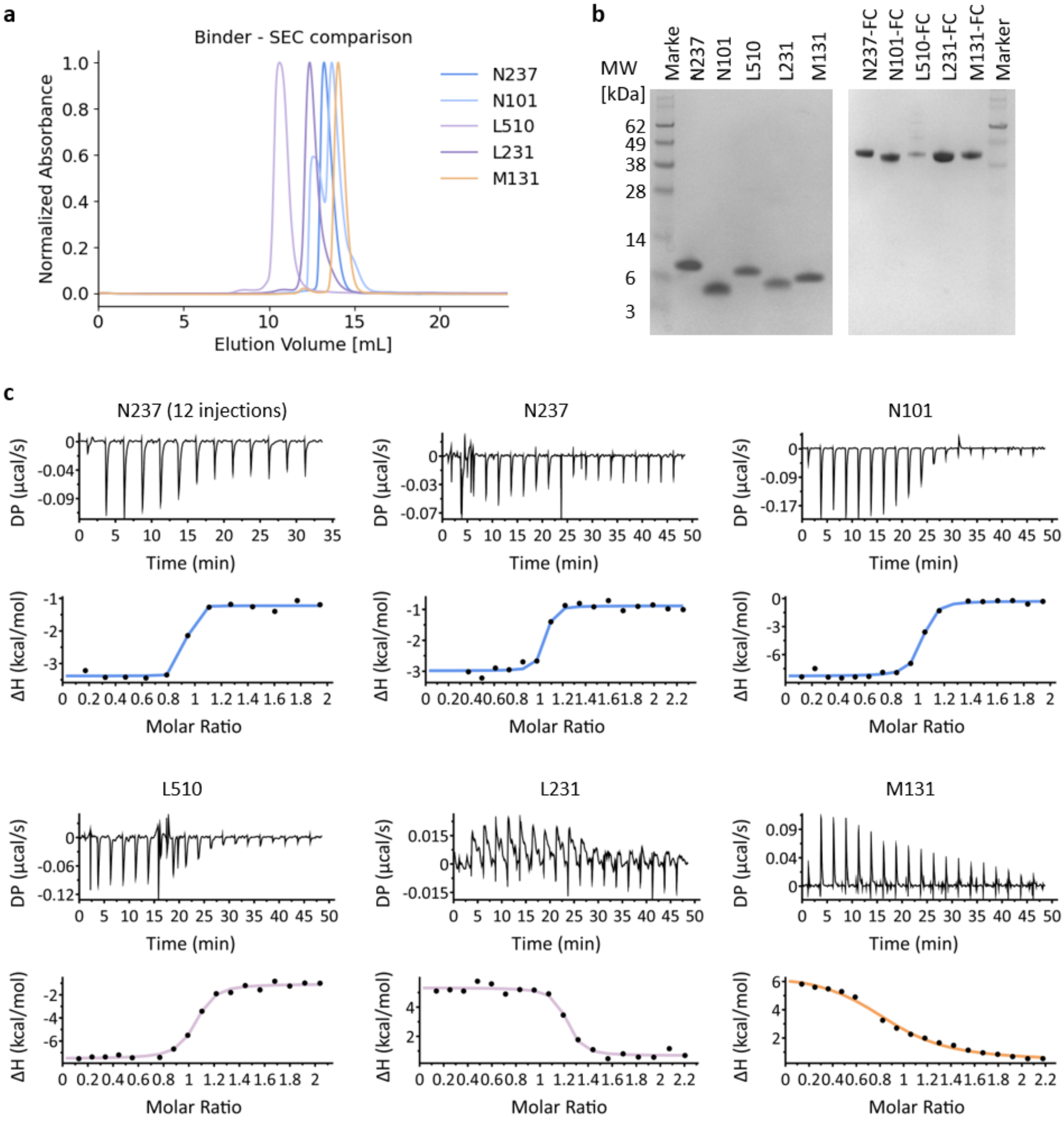


**d**

| Sample | ΔG (kcal/mol) | ΔH (kcal/mol) | -TΔS (kcal/mol) | N (sites) | $K_D$ (nM) |
|---|---|---|---|---|---|
| N237_12injections | -11.6 | -2.16 | -9.42 | 0.85 | 3.28 |
| N237 | -11.0 | -2.1 | -8.91 | 0.981 | 8.57 |
| N101 | -10.2 | -8.01 | -2.23 | 0.98 | 31.0 |
| L510 | -9.98 | -6.44 | -3.54 | 1.01 | 48.2 |
| L231 | -10.4 | 4.64 | -15.0 | 1.16 | 25.3 |
| M131 | -8.2 | 6.31 | -14.5 | 0.869 | 975.0 |

**Supplementary Fig. 4: Supplementary characterization of selected NiV, LayV and MeV RBP binders. a**, Comparison of size exclusion retention time (Superdex75 increase 10/300 GL) of selected binder without tag. Absorbance at 280 nm is normalized for each binder, respectively. **b**, SDS-PAGE of tagless and FC tagged binders. **c**, Isothermal titration calorimetry measurements of selected binder with their respective target proteins. **d**, Result table of isothermal titration calorimetry measurements.

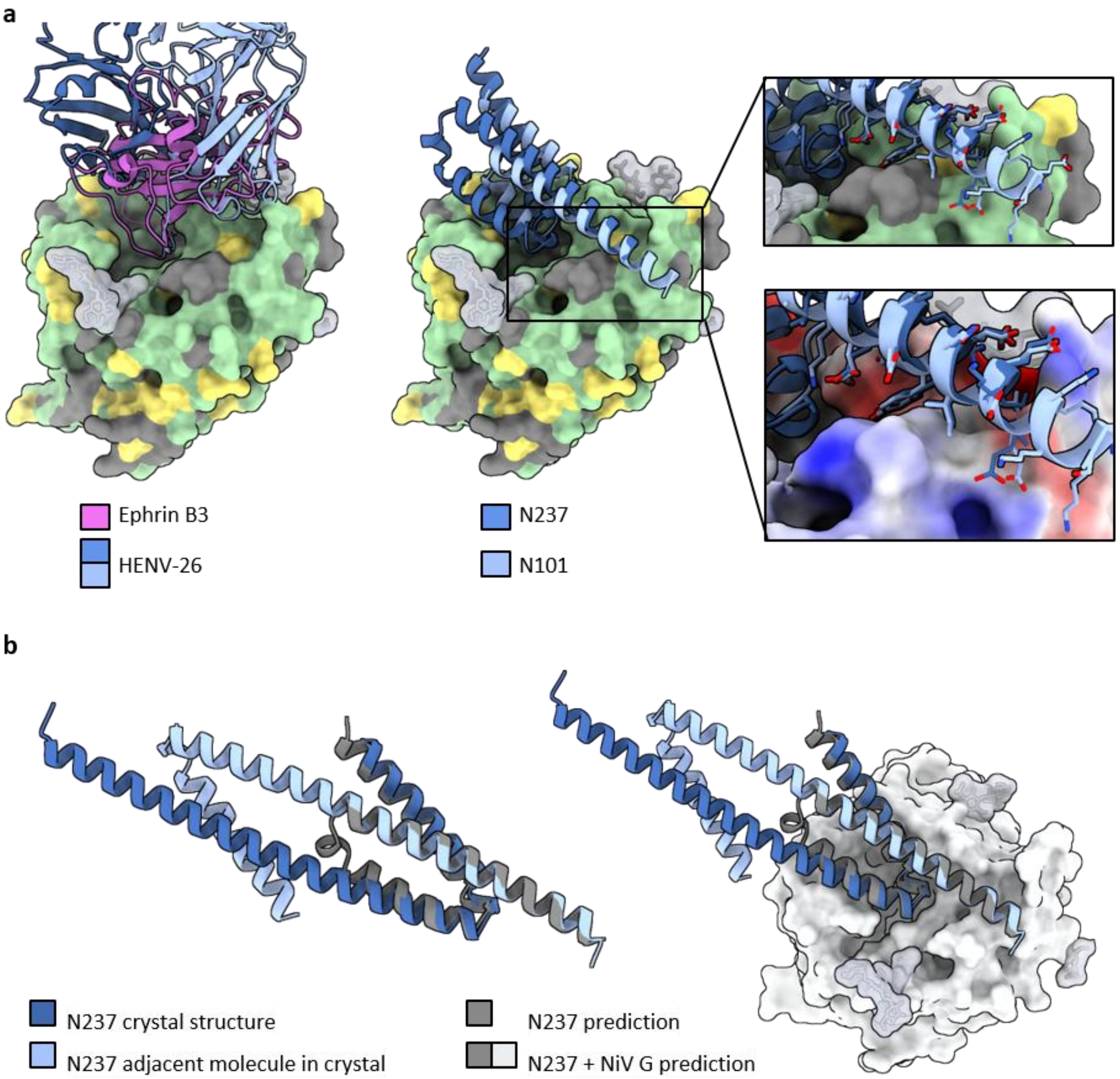


**Supplementary Fig. 5: Specificity of NiV G binders and crystal structure of N237. a**, Structure of NiV G (PDB 3D12) colored by sequence similarity to HeV G (green: identical, yellow: similar, grey: distinct). The superimposition of receptor-bound and antibody-bound NiV G (PDB 3D12, 6VY5) on the left, compared to the superimposition of N101 and N237 predictions on the right, shows that designed binders target additional residues adjacent to the receptor and antibody binding site, structurally explaining the specificity of N101 and N237 between NiV G and HeV G. **b**, Superimposition of predicted N237 and crystal structure of N237. The dimeric conformation of

N237 in the crystal structure contains the single molecule in the asymmetric unit and its neighboring molecule.

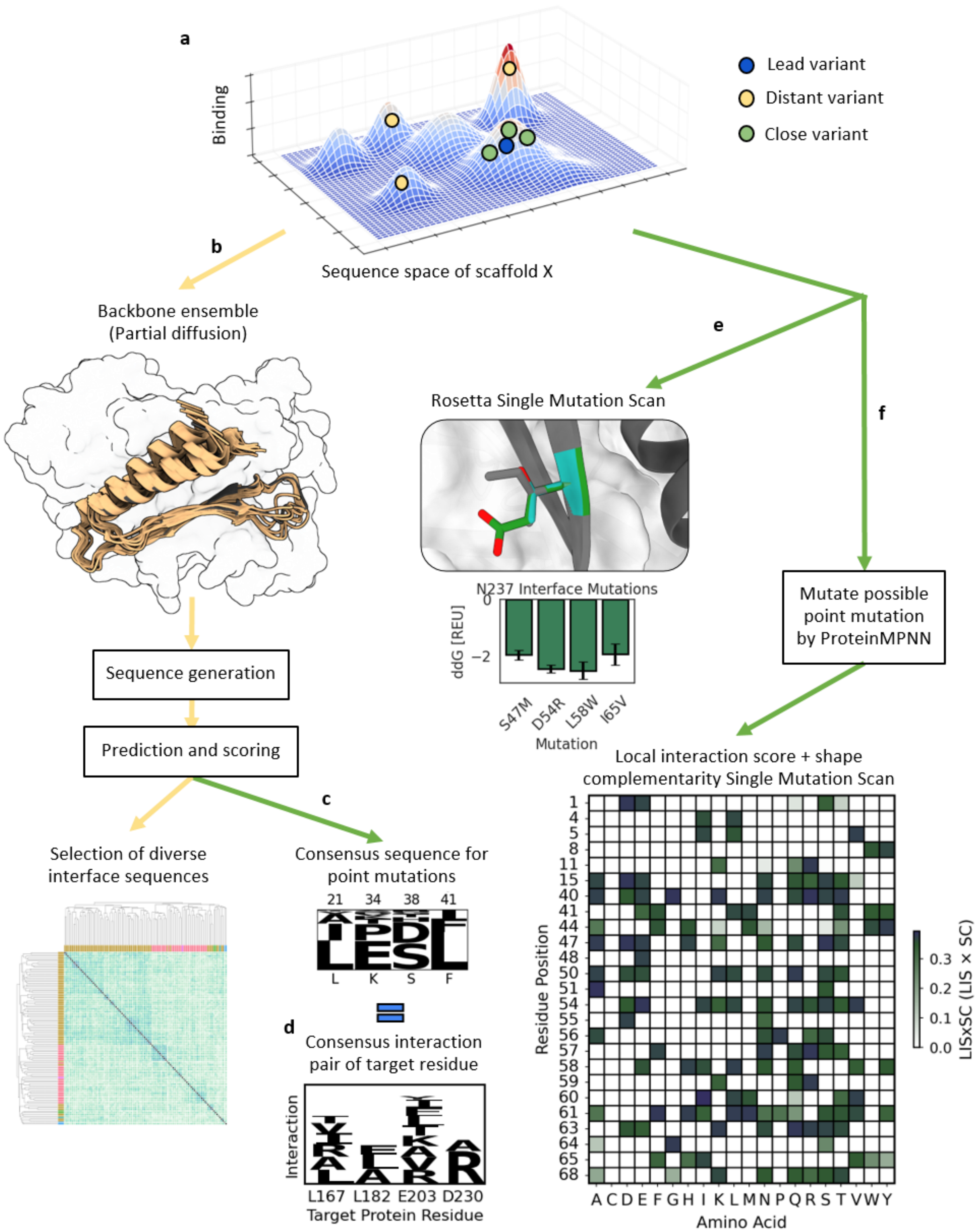
a
Binding
Sequence space of scaffold X
Lead variant
Distant variant
Close variant
b
Backbone ensemble
(Partial diffusion)
Sequence generation
Prediction and scoring
c
Selection of diverse interface sequences
Consensus sequence for point mutations
21 34 38 41
L K S F
d
Consensus interaction pair of target residue
Interaction
L167 L182 E203 D230
Target Protein Residue
e
Rosetta Single Mutation Scan
N237 Interface Mutations
ddG [REU]
0
−2
S47M D54R L58W I65V
Mutation
f
Mutate possible point mutation by ProteinMPNN
Local interaction score + shape complementarity Single Mutation Scan
Residue Position
1 4 5 8 11 15 40 41 44 47 48 50 51 54 55 56 57 58 59 60 61 63 64 65 68
A C D E F G H I K L M N P Q R S T V W Y
Amino Acid
LISxSC (LIS × SC)
0.0 0.1 0.2 0.3

**Supplementary Fig. 6: Strategies for the *in silico* optimization of binder hits. a**, The *in silico* optimization of functional binders aims to evolve the hit candidate (blue) towards a higher fitness sequence, by either introducing a few mutations (green) towards the local maxima or find another, potentially higher local maxima with a more diverse sequence (orange). **b**, A backbone ensemble is used to generate diverse sequences for a given binder scaffold. After prediction and scoring, as in the initial binder design, a diverse set of binder sequences is selected. **c**, The scored binder library is also used to find point mutations via consensus sequence, resulting in frequently sampled and well scored interface residues. **d**, The same approach, applied to an analysis of target–residue interaction partners, can be used during initial *de novo* design, providing a score that incorporates how frequently specific interactions are sampled and confidently scored. **e**, A Rosetta based mutational scan to find interface dG improving mutations was applied. **f**, Single point mutations, generated with ProteinMPNN, that improve the pAE-based local interaction score and the interface shape complementarity were determined.

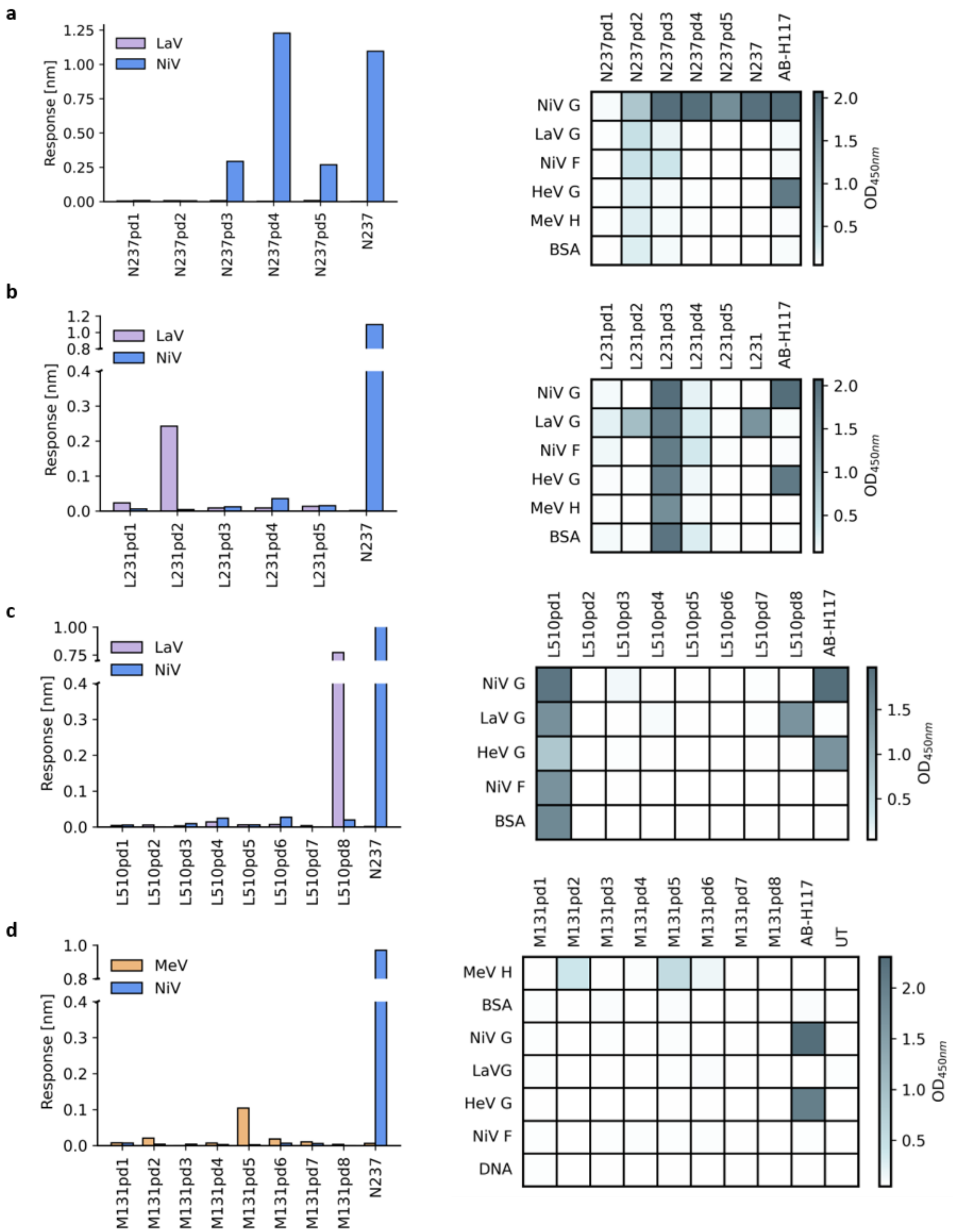
a
LaV
NiV
Response [nm]
1.25
1.00
0.75
0.50
0.25
0.00
N237pd1
N237pd2
N237pd3
N237pd4
N237pd5
N237
NiV G
LaV G
NiV F
HeV G
MeV H
BSA
AB-H117
2.0
1.5
1.0
0.5
OD450nm
b
L231pd1
L231pd2
L231pd3
L231pd4
L231pd5
L231
1.2
1.0
0.8
0.4
0.3
0.2
0.1
0.0
c
L510pd1
L510pd2
L510pd3
L510pd4
L510pd5
L510pd6
L510pd7
L510pd8
0.75
d
MeV
M131pd1
M131pd2
M131pd3
M131pd4
M131pd5
M131pd6
M131pd7
M131pd8
UT
LaVG
DNA

**Supplementary Fig. 7: Screening of partial diffused binders. a-d**, Left: BLI screening of partial diffused binders using 1000 nM of on-target and off-target protein. Right: ELISA screening using unpurified binders.

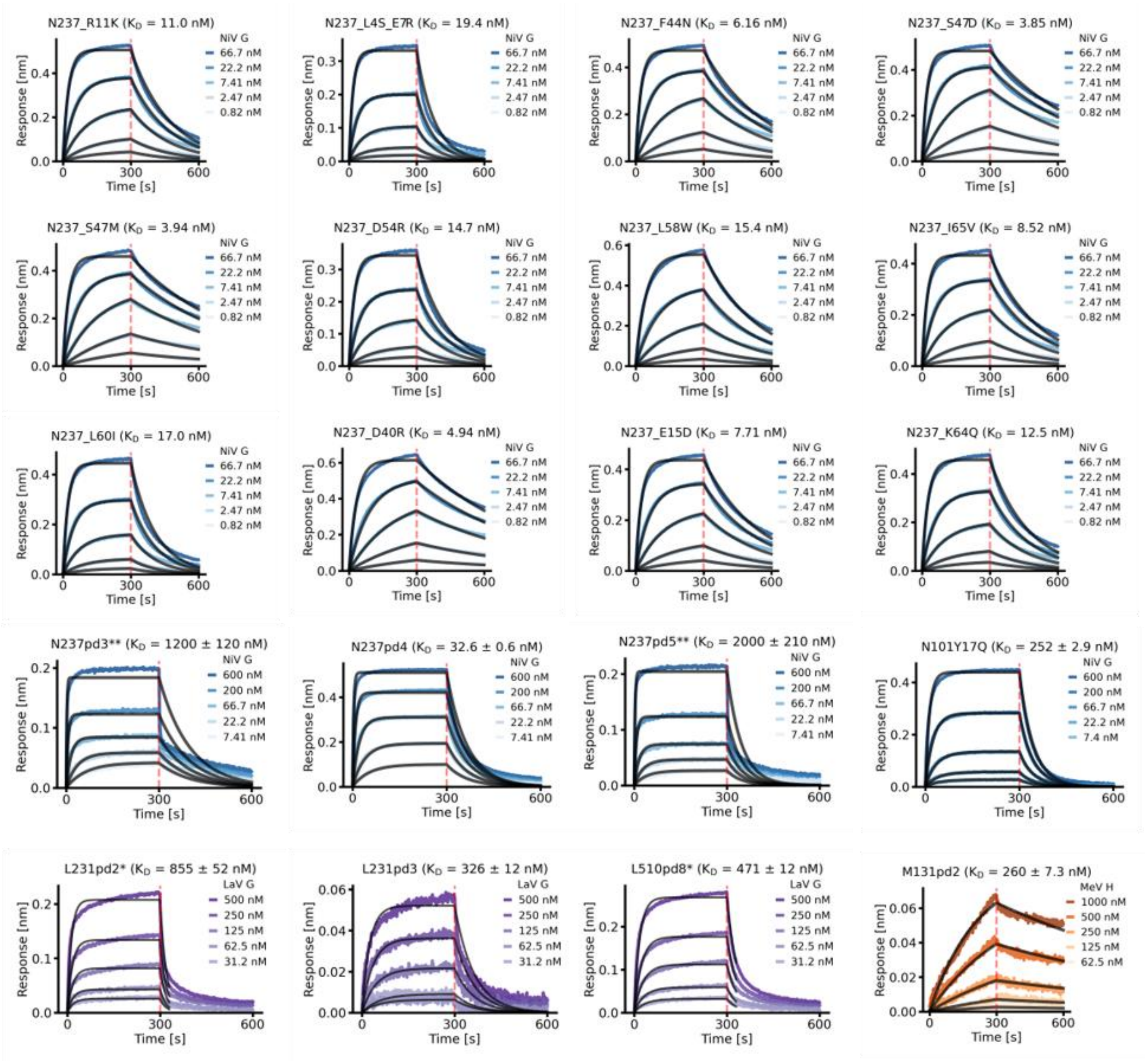


**Supplementary Fig. 8: Affinity determination of partial diffused binders and N237 point mutants.** BLI measurements of N237, N101, L231, L510, M131 variants. Each title includes the binder name and the $K_D$ of the respective measurement. * indicates reduction of the dissociation time in post-processing to improve accuracy of the fit. ** indicates $K_D$ determination via steady-state analysis.

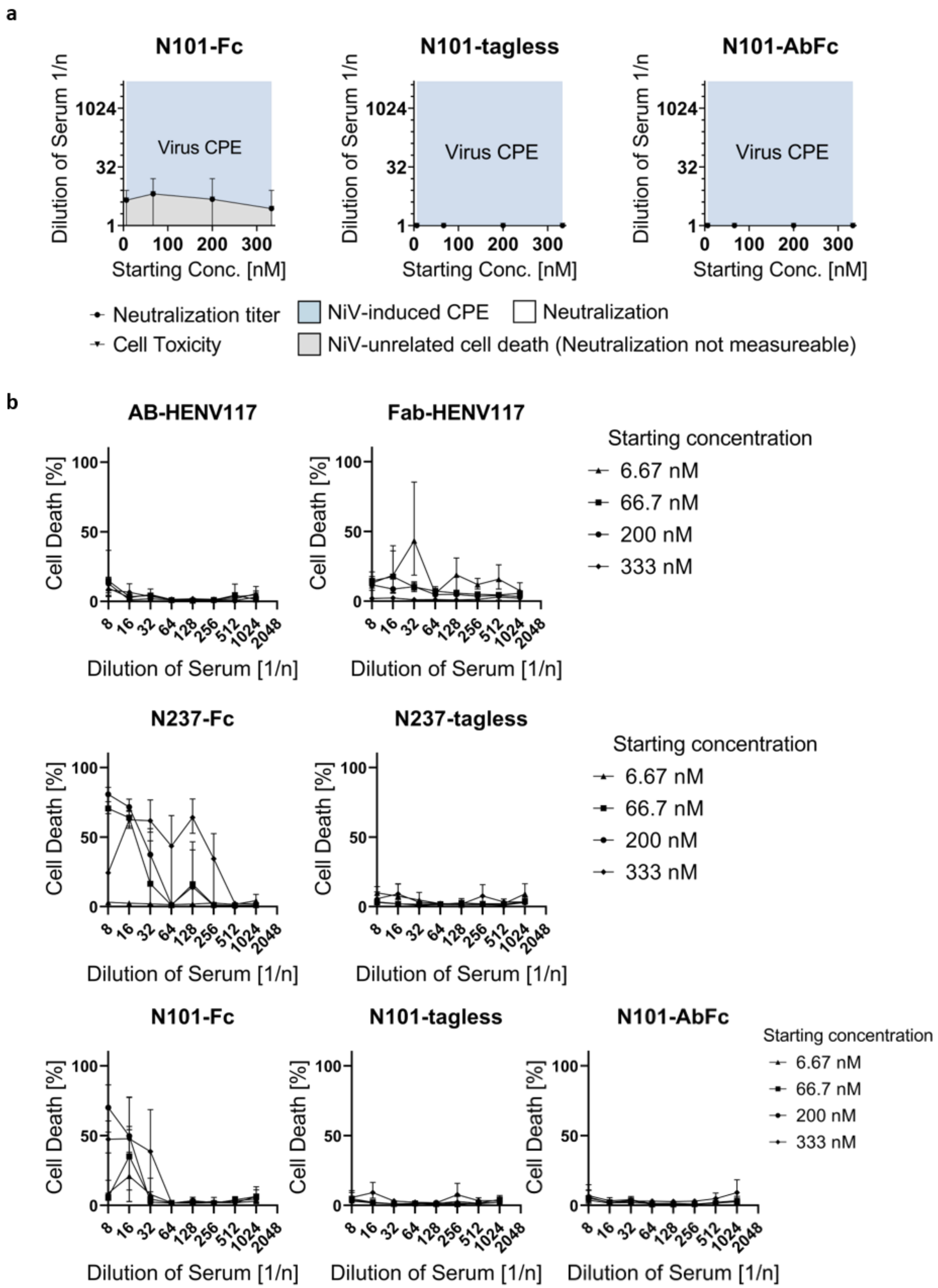
a
N101-Fc
N101-tagless
N101-AbFc
Dilution of Serum 1/n
Virus CPE
Starting Conc. [nM]
Neutralization titer
Cell Toxicity
NiV-induced CPE
NiV-unrelated cell death (Neutralization not measureable)
Neutralization
b
AB-HENV117
Fab-HENV117
N237-Fc
N237-tagless
N101-Fc
N101-tagless
N101-AbFc
Cell Death [%]
Dilution of Serum [1/n]
Starting concentration
6.67 nM
66.7 nM
200 nM
333 nM

**Supplementary Fig. 9: Neutralization and cell toxicity assay. a**, Nipah virus neutralization assay. The starting concentration of the dilution series plotted against titer of neutralization and titer of observed cell toxicity, both reported as dilution (n) of serum [1/n]. The blue area shows Nipah virus cytopathic effect (CPE), white area represents observed neutralization, grey area shows Nipah virus unrelated cell death (Cell Toxicity = Neutralization not evaluable). **b**, Determination of cell toxicity on Vero76 cells by LDH assay. Each starting concentration was diluted up to 1:2048 and cell viability/death was determined at each dilution point. LDH as cell death possible and a mock reaction were performed as controls. LDH control was set to 100% cell death.

**a**

| Target | Design | Binder Expression | Specific Binding | Unspecific Binding | Best $K_D$ (BLI) |
|---|---|---|---|---|---|
| NiV | *de novo* | 9/10 | 2/10 | 0/10 | 6.2 nM |
| | partial diffusion | 5/5 | 3/5 | 1/5 | 32.6 nM |
| LayV | *de novo* | 13/16 | 6/16 | 1/16 | 308 nM |
| | partial diffusion | 12/13 | 2/13 | 2/13 | 471 nM |
| MeV | *de novo* | 11/15 | 4/15 | 1/15 | 226 nM |
| | partial diffusion | 5/8 | 2/8 | 0/8 | 36.5 nM |
| All | | 55/67 | 19/67 | 5/67 | |

**b**

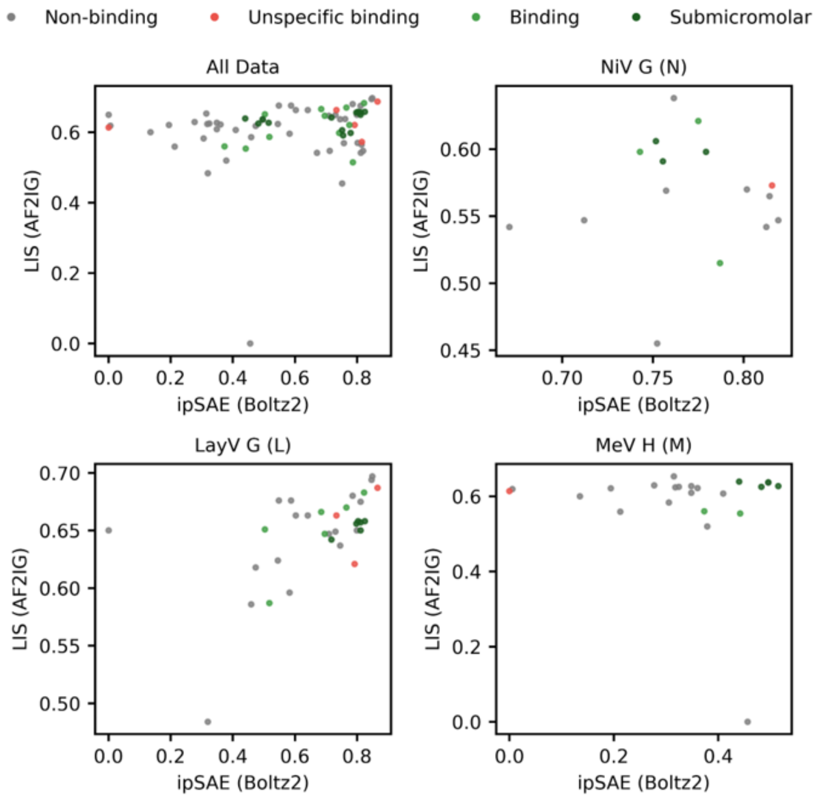


**Supplementary Fig. 10: Retroperspective AF2ig and Boltz2 analysis of experimentally characterized mini-proteins. a**, Result summary of experimental validation. **b**, Comparison of AF2ig local interaction score and Boltz2 ipSAE_min with experimental success on all three targets combined and individually.

**Supplementary Table 1. Sequences of validated binders and binder-tags.**

| Name | Sequence |
|---|---|
| C-terminal Fc-tag | GGGGSGTTCPPCPAPPVAGPSVFLFPPKPKDTLMIARTPEVTCV<br>VVDVSHEDPEVKFNWYVDGVEVHNAKTKPREEQYNSTYRVV<br>SVLTVLHQDWLNGKEYKCKVSNKALPAPIEKTISKAKGQPREP<br>QVYTLPPSRDELTKNQVSLTCLVKGFYPSDIAVEWESNGQPEN<br>NYKTTPPVLDSDGSFFLYSKLTVDKSRWQQGNVFSCSVMHEAL<br>HNHYTQKSLSLSPGKKD |
| C-terminal Ab-Fc-tag (N101-Ab-Fc) | GGGGSGDKTHTCPPCPAPELLGGPSVFLFPPKPKDTLMISRTPE<br>VTCVVVDVSHEDPEVKFNWYVDGVEVHNAKTKPREEQYNST<br>YRVVSVLTVLHQDWLNGKEYKCKVSNKALPAPIEKTISKAKG<br>QPREPQVYTLPPSREEMTKNQVSLTCLVKGFYPSDIAVEWESN<br>GQPENNYKTTPPVLDSDGSFFLYSKLTVDKSRWQQGNVFSCSV<br>MHEALHNHYTQKSLSLSPGK |
| C-terminal Fc-tag (M131pd-variants) | GGGGSGGGGSGTTCPPCPAPPVAGPSVFLFPPKPKDTLMIARTP<br>EVTCVVVDVSHEDPEVKFNWYVDGVEVHNAKTKPREEQYNST<br>YRVVSVLTVLHQDWLNGKEYKCKVSNKALPAPIEKTISKAKG<br>QPREPQVYTLPPSRDELTKNQVSLTCLVKGFYPSDIAVEWESNG<br>QPENNYKTTPPVLDSDGSFFLYSKLTVDKSRWQQGNVFSCSVM<br>HEALHNHYTQKSLSLSPGKKD |
| N-terminal HIS-TEV-tag | MGHHHHHHGGENLYFQG |

| | |
|---|---|
| N237 | EKELIEEYEKREKEELEELEKQIKELKELIDKLGKEKYKDLYTFLESEFAAAKEDPRLRLLLKGIIKKKIAELKAELAAKEA |
| N101 | SEKKKREEELIKEYEEYEKKKAELLKEITALDPALADFAVYSNLITEELLAKAKARAAAAAA |
| L231 | LPPMPTPEEIRAKIRELMNKVAELLKGHEEEVNAIIKKYVDNGLYDEAGTEIMRIAIEAKNEEALKALAEVGRALREIFDEVRARAAARR |
| L510 | GPSPEEKAQLLKENMYLIVDRTGELYERLKEANVPRRYRAALAFTAETLLVDYLHMAVENNKVVSVEELEEYWETEVRAWVERMLAILQKNFPEIKPIKW |
| M131 | MTELEKAEKLAKELEKRLEELGAKNVKVTIEENKEDTSGLFARKSAIVKYEIPDSATPEQKSRIMSALTSGLRKIATALGKAE |
| M131pd5 | MSAEEELEKLAEEAKKELEKLGAEDVEVKVIENKDRADPALGERSVVITYKIPESFSPEEKSKIMAALEVNMRKLATALLKAE |

**Supplementary Table 2. Sequences of viral glycoproteins, antibodies and ephrin B2.**

| Name | Sequence |
|---|---|
| NiV G | EGVSNLVGLPNNICLQKTSNQILKPKLISYTLPVVGQSGTCITDP<br>LLAMDEGYFAYSHLERIGSCSRGVSKQRIIGVGEVLDRGDEVPS<br>LFMTNVWTPPNPNTVYHCSAVYNNEFYYVLCAVSTVGDPILNS<br>TYWSGSLMMTRLAYVKPKSNGGGYNQHQLALRSIEKGRYDK<br>VMPYGPSGIKQGDTLYFPAVGFLVRTEFKYNDSNCPITKCQYS<br>KPENCRLSMGIRPNSHYILRSGLLKYNLSDGENPKVVFIEISDQR<br>LSIGSPSKIYDSLGQPVFYQASFSWDTMIKFGDVLTVNPLVVNW<br>RNNTVISRPGQSQCPRFNTCPEICWEGVYNDAFLIDRINWISAG<br>VFLDSNQTAENPVFTVFKDNEILYRAQLASEDTNAQKTITNCFL<br>LKNKIWCISLVEIYDTGDNVIRPKLFAVKIPEQCYGGSGLNDIFE<br>AQKIEWHEGSGGHHHHHH |
| NiV F | ILHYEKLSKIGLVKGVTRKYKIKSNPLTKDIVIKMIPNVSNMSQC<br>TGSVMENYKTRLNGILTPIKGALEIYKNNTHDLVGDVRLAGVI<br>MAGVAIGIATAAQITAGVALYEAMKNADNINKLKSSIESTNEA<br>VVKLQETAEKTVYVLTALQDYINTNLVPTIDKISCKQTELSLDL<br>ALSKYLSDLLFVFGPNLQDPVSNSMTIQAISQAFGGNYETLLRT<br>LGYATEDFDDLLESDSITGQIIYVDLSSYYIIVRVYFPILTEIQQA<br>YIQELLPVSFNNDNSEWISIVPNFILVRNTLISNIEIGFCLITKRSVI<br>CNQDYATPMTNNMRECLTGSTEKCPRELVVSSHVPRFALSNGV<br>LFANCISVTCQCQTTGRAISQSGEQTLLMIDNTTCPTAVLGNVII<br>SLGKYLGSVNYNSEGIAIGPPVFTDKVDISSQISSMNQSLQQSKD |

| | |
|---|---|
| | YIKEAQRLLDTVNPSMKQIEDKIEEILSKIYHIENEIARIKKLIGE<br>APGGTKHHHHHH |
| LayV G | TIKPVEYYKPDGCNKTNDHFTMQPGVNFYTVPNLGPSSSSADE<br>CYTNPSFSIGSSIYMFSQEIRKTDCTTGEILSIQIVLGRIVDKGQQ<br>GPQASPLLVWSVPNPKIINSCAVAAGDETGWVLCSVTLTAASG<br>EPIPHMFDGFWLYKFEPDTEVVAYRITGFAYLLDKVYDSVFIGK<br>GGGIQRGNDLYFQMFGLSRNRQSIKALCEHGSCLGTGGGGYQ<br>VLCDRAVMSFGSEESLISNAYLKVNDVASGKPTIISQTFPPSDSY<br>KGSNGRIYTIGERYGIYLAPSSWNRYLRFGLTPDISVRSTTWLK<br>EKDPIMKVLTTCTNTDKDMCPEICNTRGYQDIFPLSEDSSFYTYI<br>GITPSNEGTKSFVAVKDDAGHVASITILPNYYSITSATISCFMYK<br>EEIWCIAVTEGRKQKENPQRIYAHSYRVQKMCFNIKPAGGSGL<br>NDIFEAQKIEWHEGSGGHHHHHH |
| HeV G | QGVSDLVGLPNQICLQKTTSTILKPRLISYTLPINTREGVCITDPL<br>LAYVDNGFFAYSHLEKIGSCTRGIAKQRIIGVGEVLDRGDKVPS<br>MFMTNVWTPPNPSTIHHCSSTYHEDFYYTLCAVSHVGDPILNST<br>SWTESLSLIRLAYVRPKSDSGDYNQKYIAITKVERGKYDKVMP<br>YGPSGIKQGDTLYFPAVGFLPRTEFQYNDSNCPIIHCKYSKAEN<br>CRLSMGVNSKSHYILRSGLLKYNLSLGGDIILQFIEIADNRLTIGS<br>PSKIYNSLGQPVFYQASYSWDTMIKLGDVDTVDPLRVQWRNN<br>SVISRPGQSQCPRFNVCPEVCWEGTYNDAFLIDRLNWVSAGVY<br>LNSNQTAENPVFAVFKDNEILYQVPLAEDDTNAQKTITDCFLLE |

| | |
|---|---|
| | NVIWCISLVEIYDTGDSVIRPKLFAVKIPAQCSGGSGLNDIFEAQ KIEWHEGSGGHHHHHH |
| MeV H | ADVAAEELMNALVNSTLLETRTTNQFLAYVSKGNCSGPTTIRG QFSNMSLSLLDLYLGRGYNVSSIVTMTSQGMYGGTYLVEKPNL SSKRSELSQLSMYRVFEVGVIRNPGLGSPVFHMTNYLEQPVSN DLSNCMVALGELKLAALCHGEDSITIPYQGSGKGVNFQLVKLG VWKSPTDMQSWVPLSTDDPVIDRLYLSSHRGVIADNQAKWAV PTTRTDDKLRMETCFQQACKGKIQALCENPEWAPLKDNRIPSY GVLSVDLSLTVELKIKIASGFGPLITHGSGMDLYKSNHNNVYW LTIPPMKNLALGVINTLEWIPRFKVSPYLFTVPIKEAGEDCHAPT YLPAEVDGDVKLSSNLVILPGQDLQYVLATYDTSRVEHAVVY YVYSPSRSFSYFYPFRLPIKGVPIELQVECFTWDQKLWCRHFCV LADSESGGHITHSGMVGMGVSCTVTGGSGLNDIFEAQKIEWHE GSGGHHHHHH |
| HENV-117-HC | QVQLQQWGAGLLKPSETLSLTCDLSGGSFTNYYWTWIRHLPG KGLEWLGEINHSGRTNYSPSLKSRVTISVDTSKNQFSLRLSSVT AADTAVYYCARDIRKLAPKPHSYDLYYGMDVWGQGTTVTVS S |
| HENV-117-LC | EIVLTQSPGTLSLSPGERATLSCRASQSVGSNYLTWYQQKPGQP PRLLIYGASNRAAGIPDRFSGSGSGTDFTLTISRLEPEDFAVYYC QQYETSSRTFGQGTKVDIR |

| | |
|---|---|
| HENV-32-HC | QVQLVQSGAEVKKPGSSVKVSCKVSGGIFNRETINWVRQAPGQGLEWMGRITPIVDVPNYPRKFRGRVTITADKSTSTVYMELSGLRFEDTAIYFCARFRGHNYFDPWGQGTLVTVSS |
| HENV-32-LC | SYVLTQPPSVSVAPGQTARITCGGNNIGGKSVHWYQQKPGQAPVLVVYDDRDRPSGIPERFSGSNSGDTASLTISRVDAGDEADYFCQVWDNASDEAVFGGGTKLT |
| Ephrin B2 | CLSFWQELRPSQAEFKLATMGILPSPGMPALLSLVSLLSVLLMGCVAETGSIVLEPIYWNSSNSKFLPGQGLVLYPQIGDKLDIICPKVDSKTVGQYEYYKVYMVDKDQADRCTIKKENTPLLNCAKPDQDIKFTIKFQEFSPNLWGLEFQKNKDYYIISTSNGSLEGLDNQEGGVCQTRAMKILMKVGQDGTKHHHHHH |
| NiV G (Ephrin B2 competition assay) | GLPNNICLQKTSNQILKPKLISYTLPVVGQSGTCITDPLLAMDEGYFAYSHLERIGSCSRGVSKQRIIGVGEVLDRGDEVPSLFMTNVWTPPNPNTVYHCSAVYNNEFYYVLCAVSTVGDPILNSTYWSGSLMMTRLAYVKPKSNGGGYNQHQLALRSIEKGRYDKVMPYGPSGIKQGDTLYFPAVGFLVRTEFKYNDSNCPITKCQYSKPENCRLSMGIRPNSHYILRSGLLKYNLSDGENPKVVFIEISDQRLSIGSPSKIYDSLGQPVFYQASFSWDTMIKFGDVLTVNPLVVNWRNNTVISRPGQSQCPRFNTCPEICWEGVYNDAFLIDRINWISAGVFLDSNQTAENPVFTVFKDNEILYRAQLASEDTNAQKTITNCFLLKNKIWCISLVEIYDTGDNVIRPKLFAVKIPEQCT |

**Supplementary Table 3. Crystallographic data collection and refinement statistics**.

| | L510 | N237 |
|---|---|---|
| **PDB ID** | 9TNN | 9TNU |
| **Data collection**[a] | | |
| Beamline | BESSY MX14.2 | EMBL PETRA III P14 |
| Resolution range (Å) | 41.7 – 1.93 (2.05 – 1.93) | 46.9 – 1.39 (1.59 – 1.39) |
| Space group | C 2 2 21 | P 31 2 1 |
| Unit cell dimensions | | |
| a, b, c (Å) | 61.81 109.51 64.33 | 54.21 54.21 59.94 |
| α, β, γ (°) | 90 90 90 | 90 90 120 |
| Observed reflections | 208285 (24752) | 194833 (10813) |
| Unique reflections | 16674 (2557) | 5935 (541) |
| Rmerge | 0.34 (1.73) | 0.07 (1.15) |
| Rmeas | 0.35 (1.82) | 0.07 (1.20) |
| CC1/2 | 0.995 (0.468) | 0.999 (0.738) |
| Multiplicity | 12.49 (9.68) | 18.0 (11.0) |
| <I/σI> | 8.20 (1.15) | 17.3 (1.8) |
| Completeness (%) | | |
| Spherical | 99.3 (95.7) | 51.6 (8.0) |
| Ellipsoidal | | 91.2 (72.2) |
| **Refinement** | | |
| Protein molecules/ASU | 2 | 1 |
| Resolution range (Å) | 41.7 – 1.93 (2.05 – 1.93) | 46.9 - 1.89 (2.16 - 1.89) |
| Reflections all/free R set | 16668 (2633) / 832 (132) | 7978 (2508) / 398 (130) |
| $R_{work}/R_{free}$ | 0.218/0.239 | 0.238 / 0.256 |
| No. of atoms: | | |
| Protein | 1701 | 636 |
| Water | 166 | 121 |
| Non-water solvent | 28 | 8 |
| Mean B-factors (Å$^2$): | | |
| Protein | 31.04 | 25.31 |
| Water | 39.79 | 38.15 |
| Non-water solvent | 39.34 | 41.06 |
| RMSD: | | |
| Bond lengths (Å) | 0.017 | 0.017 |
| Bond angles (°) | 1.57 | 1.47 |
| Ramachandran statistics | | |
| Favored/allowed/outliers (%) | 99.49 / 0.51 / 0 | 96.20 / 2.53 / 1.27 |

[a]Numbers in parentheses correspond to the highest resolution shells.
